# Towards a phylogenomic classification of Magnoliidae

**DOI:** 10.1101/2024.01.09.574948

**Authors:** Andrew J. Helmstetter, Zacky Ezedin, Elton John de Lírio, Sylvia M. de Oliveira, Lars W. Chatrou, Roy H.J. Erkens, Isabel Larridon, Kevin Leempoel, Olivier Maurin, Shyamali Roy, Alexandre R. Zuntini, William J. Baker, Thomas L.P. Couvreur, Félix Forest, Hervé Sauquet

**Author notes:** Joint senior authors (William J. Baker, Thomas L.P. Couvreur, Félix Forest).

## Abstract

**Premise:** Magnoliidae are a strongly supported clade of angiosperms. Previous phylogenetic studies based primarily on analyses of a limited number of mostly plastid markers have led to the current classification of magnoliids into four orders and 18 families. However, uncertainty remains regarding the placement of several families.

**Methods:** Here we present the first comprehensive phylogenomic analysis of Magnoliidae as a whole, sampling 235 species from 199 (74%) genera and representing all families and most previously accepted subfamilies and tribes. We analyze newly generated data from the Angiosperms353 probe set using both coalescent and concatenation analyses and testing the impact of multiple filtering and alignment strategies.

**Results:** While our results generally provide further support for previously established phylogenetic relationships in both magnoliids as a whole and large families including Annonaceae and Lauraceae, they also provide new evidence for previously ambiguous relationships. In particular, we find support for the position of Hydnoraceae as sister to the remainder of Piperales and, for the first time, resolve the backbone of relationships among most genera of Myristicaceae.

**Conclusions:** Although some of our results are limited by low gene recovery for a number of taxa and significant gene tree conflict for some relationships, this study represents a significant step towards reconstructing the evolutionary history of a major lineage of angiosperms. Based on these results, we present an updated phylogenetic classification for Magnoliidae, recognizing 21 families, summarizing previously established subfamilies and tribes, and describing new tribes for Myristicaceae.

## INTRODUCTION

Magnoliidae (sensu Cantino et al., 2007) are a large clade of angiosperms consisting of approximately 10,000 species assigned to 18 families and four orders (Massoni et al., 2014; APG IV, 2016; Stevens, 2022). Magnoliidae have evolved separately from other major lineages of angiosperms (incl. monocots, eudicots, *Amborella*, Nymphaeales, Austrobaileyales, Chloranthaceae, and *Ceratophyllum*) for a long time, as exemplified by their presence in the fossil record since at least the Barremian (ca. 121-129 Ma; Massoni et al., 2015). Species of Magnoliidae are found across a broad range of habitats and climates, but occur predominantly in tropical and warm temperate rain forests. The group has long been noted for its diverse and unusual flower morphology (Figs. 1, 2; Endress, 1987), leading to speculations that some living members of the group may resemble ancestors of angiosperms as a whole (Cronquist, 1981). Although many questions remain, phylogenetic analyses of morphological evolution have consistently rejected this idea (Sauquet et al., 2003, 2017; Endress and Doyle, 2009).

**FIGURE 1.**
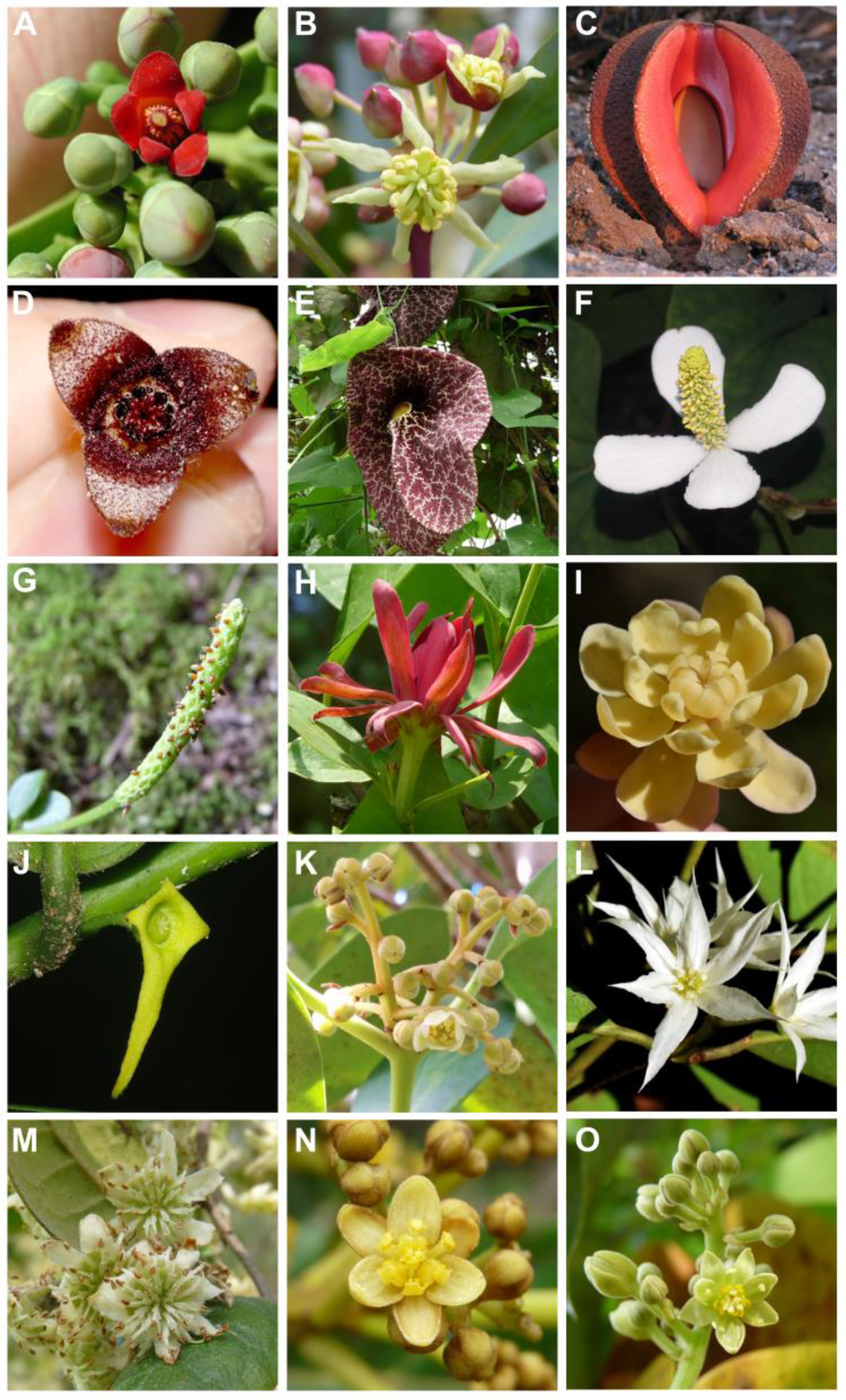
Floral diversity in Canellales, Piperales, and Laurales. A) Canellaceae: *Canella winterana* (L.) Gaertn. (Kevin Nixon, asked for permission); B) Winteraceae, Winteroideae: *Drimys lanceolata* (Poir.) Baill.; C) Hydnoraceae, *Hydnora visseri* Bolin, E.Maass & Musselman; D) Asaraceae: *Asarum europaeum* L.; E) Aristolochiaceae: *Aristolochia gigantea* Mart. & Zucc.; F) Saururaceae: *Houttuynia cordata* Thunb.; G) Piperaceae, Piperoideae: *Peperomia tetraphylla* (G.Forst.) Hook. & Arn.; H) Calycanthaceae, Calycanthoideae: *Calycanthus occidentalis* Hook. & Arn.; I) Calycanthaceae, Idiospermoideae: *Idiospermum australiense* (Diels) S.T.Blake; J) Siparunaceae: *Glossocalyx longicuspis* Benth.; K) Gomortegaceae: *Gomortega keule* (Molina) Baill.; L) Atherospermataceae: *Doryphora sassafras* Endl.; M) Monimiaceae, Monimioideae: *Peumus boldus* Molina; N) Lauraceae, Hypodaphnideae: *Hypodaphnis zenkeri* (Engl.) Stapf; O) Lauraceae, Laureae: *Persea americana* L. Photo credits: A: Kevin Nixon (http://www.diversityoflife.org/); C: CC BY https://commons.wikimedia.org/wiki/File:Hydnora_visseri,_Namibia.png; B, D, E, G, H, I, L, M, N, O: Hervé Sauquet; F: Laetitia Carrive; J: Thomas L.P. Couvreur; K: Ana Almeida.

Numerous studies at various scales conducted over the last three decades have led to an increasingly robust and well supported backbone for phylogenetic relationships among families of Magnoliidae (Qiu et al., 1999; Doyle and Endress, 2000; Sauquet et al., 2003; Moore et al., 2007; Soltis et al., 2011; Massoni et al., 2014; Wickett et al., 2014; Li et al., 2019; One Thousand Plant Transcriptomes Initiative, 2019). Briefly, these studies have supported the monophyly of the four orders as initially recircumscribed by the Angiosperm Phylogeny Group (APG, 1998; APG II, 2003), with two sister clades, one comprising Canellales and Piperales, and the other Laurales and Magnoliales. Relationships within orders have become relatively stable as well, but important questions persist. In particular, much uncertainty remains in Piperales regarding the relationship between the two traditional lineages of Aristolochiaceae (subfamilies Asaroideae and Aristolochioideae) and two enigmatic taxa (subfamily Hydnoroideae and *Lactoris*) (Naumann et al., 2013; Massoni et al., 2014; Jost et al., 2021). In Laurales, no consensus has emerged yet on the relationships among the three largest families, Lauraceae, Hernandiaceae, and Monimiaceae (Doyle and Endress, 2000; Renner and Chanderbali, 2000; Massoni et al., 2014). In Magnoliales, the position of Magnoliaceae in relation to the other families of the order also continues to be equivocal (Doyle and Endress, 2000; Sauquet et al., 2003; Massoni et al., 2014). Lastly, considerable progress has been made in reconstructing the phylogenetic backbones of two of the largest families, Annonaceae (Chatrou et al., 2012; Guo et al., 2017; Couvreur et al., 2019) and Piperaceae (Jaramillo et al., 2004; Wanke et al., 2007b, a; Smith et al., 2008). In contrast, the global phylogeny of Lauraceae is only now becoming more confidently resolved (Song et al., 2020) and very few studies have addressed relationships among genera in two smaller families, Monimiaceae (Renner et al., 2010) and Myristicaceae (Sauquet et al., 2003; Massoni et al., 2014).

Here we take advantage of recent technological advances in plant nuclear phylogenomics to further explore phylogenetic relationships among genera and families of magnoliids. Specifically, we use a target enrichment strategy with the universal Angiosperms353 probe set (Johnson et al., 2019; Baker et al., 2022). Although a specific probe set was developed for Annonaceae and may be effective in other magnoliid families (Couvreur et al., 2019), there is growing evidence that universal probes may be as effective as clade-specific probes (Larridon et al., 2020; Shah et al., 2021) and the benefits of community convergence around a standard gene set are well-documented (Baker et al., 2021; McDonnell et al., 2021). However, the utility of Angiosperms353, and targeted sequence capture in general, to resolve relationships in a clade that contains both very deep and very shallow divergence events remains a salient question. Our goal here was to capitalize on the unique opportunity created by the extensive sequencing of magnoliid genera undertaken jointly by the Plant and Fungal Trees of Life (PAFTOL; Baker et al., 2022) project and the Genomics for Australian Plants (GAP) project, in order to evaluate the utility of these new data to further test and address long-standing questions remaining about the phylogenetic backbone of Magnoliidae. In doing so, we also aimed at testing the impact of multiple data filtering strategies on phylogenomic reconstruction and associated measures of branch support. While our results broadly corroborate with greater support most of the relationships that had emerged previously, we also find new evidence for the positions of Lauraceae and Magnoliaceae and a distinct placement for Hydnoraceae previously not retrieved. We also resolved, for the first time, most relationships among genera in the enigmatic Myristicaceae. Based on this new study and previous work in selected families, we propose an updated phylogenetic classification for Magnoliidae as a whole.

## MATERIALS AND METHODS

### Taxon sampling

Our sampling consists of 238 taxa representing 235 unique species, 199 genera, and all 18 families of Canellales, Laurales, Magnoliales, and Piperales, with a respective generic representation of 90% (9/10), 71% (75/106), 74% (101/136) and 88% (14/16) according to the revised list of genera presented at the end of this paper. In addition, we used six specimens sampled from the four genera of Chloranthaceae (Chloranthales) as outgroup taxa in phylogenetic analyses, a choice supported by most recent phylogenomic analyses of nuclear data (One Thousand Plant Transcriptomes Initiative, 2019; Baker et al., 2022). Samples for this study were obtained from five distinct sources. The vast majority (191) were new sequence data generated for this study, as part of the broader PAFTOL project.

Another 20 taxa from Australia were new sequence data contributed by the GAP project, and six additional taxa were sourced from a project on Neotropical Myristicaceae led by one of us (SMDO). Lastly, we also used previously published sequence data for 22 taxa from the One Thousand Plant Transcriptomes (1KP) Initiative (2019) and five from the Sequence Read Archive (SRA). All voucher and source details are provided as Appendix S1.

### DNA extraction, library preparation and sequencing

For PAFTOL samples, we extracted DNA using a modified CTAB protocol (Doyle and Doyle, 1987) and then purified it using Mag-Bind® TotalPure NGS magnetic beads (Omega Bio-tek, Norcross, Georgia, USA). We then assessed the average fragment size within each extraction using 1.5x agarose gel and quantified the amount of double stranded DNA using a Qubit® 3.0 fluorometer (ThermoFisher Scientific, Waltham, Massachusetts, USA). If average fragment size was longer than 350 base pairs (bp), extracts were sonicated with a M220 Focused-ultrasonicator™ with microTUBEs AFA Fiber Pre-Slit Snap-Cap (Covaris, Woburn, Massachusetts, USA) following the manufacturer’s protocol to generate ∼350 bp insert sizes.

We prepared dual-indexed libraries with Dual Index Primers Set 1, NEBNext® Multiplex Oligos for Illumina® (New England BioLabs, Ipswich, Massachusetts, USA) using DNA NEBNext® UltraTM II Library Prep Kits at half the recommended volume. We examined the quality of each library using a 4200 TapeStation and High Sensitivity D1000 ScreenTapes (Agilent Technologies, Santa Clara, California, USA). Again, we quantified the amount of DNA with a Qubit fluorometer. Libraries were pooled so that they were equimolar and contained 20 to 25 individual libraries totaling 1 μg of DNA. These pools were hybridized using the myBaits Expert Predesigned Panel (Arbor Biosciences, Ann Arbor, Michigan, USA) Angiosperms353 v1 (Catalog #308196) (Johnson et al., 2019) following the manufacturer’s protocol with v4 chemistry (http://www.arborbiosci.com/mybaits-manual). We hybridized libraries at 65°C for up to 32 hours in a Hybex™ Microsample Incubator. To prevent evaporation, we added an equal volume of red Chill-out™ Liquid Wax (Bio-Rad, Hercules, California, USA) to each hybridization reaction.

We amplified the enriched products using the KAPA HiFi (2x) HotStart ReadyMix PCR Kit (Roche, Basel, Switzerland) for 10 cycles. PCR products were then cleaned using the QIAquick PCR purification kit (Qiagen). We performed another quantification with the Qubit fluorometer and, if necessary, re-amplified enriched products a second time for 3–8 cycles. We assessed quality and fragment size a final time using the TapeStation as above. Enriched library pools were then multiplexed and sequenced on an Illumina MiSeq (Illumina, San Diego, California, USA) with v3 reagent chemistry (2 × 300 bp paired-end reads) at Royal Botanic Gardens, Kew, or on an Illumina HiSeq (2 × 150 bp paired-end reads) at Genewiz (Takeley, UK) or at Macrogen (Geumcheon, Republic of Korea).

For GAP samples, dried plant tissue (20–30 mg) was ground using a TissueLyser II (Qiagen) with tungsten carbide beads for simultaneous disruption and homogenization of the sample, per the manufacturer’s instructions. Genomic DNA was extracted using the DNeasy® Plant mini kit (Qiagen) as per the manufacturer’s instructions on a QIAcube Connect (Qiagen). DNA quantity and quality was assessed using 1% E-gel with Sybr Safe dye (ThermoFisher) and concentrations assessed using Quantifluor dsDNA assay (Promega). DNA samples were fragmented enzymatically as part of the NEBNext Ultra II FS library preparation workflow.

Libraries were prepared using the NEBNext Ultra II FS Library Prep Kit (New England Biolabs, Ipswich, MA, USA), following the manufacturer’s instructions with inserts of approximately 350 bp. Capture pools were 12–16 plex. Pooled libraries were enriched using the Angiosperms353 probe kit (Johnson et al. 2018) by hybridizing at 65°C with the Arbor Biosciences MyBaits Expert Plant Angiosperms353 v1 probe set with V5 chemistry (Cat. # 308108.v5) following the manufacturer’s instructions. Sequencing was performed on a NovaSeq 6000 (Illumina Inc., San Diego, USA) at the Australian Genome Research Facility (Melbourne) with v1.5 chemistry and 150bp paired-end reads.

### Contig assembly, multi-sequence alignment and paralog detection

We used HybPiper (v1.2) (Johnson et al., 2016) with default settings to process the raw sequence data. Briefly, reads were mapped to targets using BWA (v0.7.5a) (Li and Durbin, 2009) and successfully mapped reads were assembled into contigs using SPAdes (v3.11.1) (Bankevich et al., 2012). Then, Exonerate (Slater and Birney, 2005) was used to align contigs to their associated target exon sequence. If contigs were found to be overlapping (Johnson et al., 2016) they were combined into “supercontigs” that contain both target (exon) and off-target sequence data. Exonerate was then run again so that off-target regions could be more accurately identified.

Multi-sequence alignments were generated using two methods: MAFFT (v7.305) (Katoh and Standley, 2013) using the “auto” option and MAGUS (Smirnov and Warnow, 2021). The former aligns using Fourier transforms and automatically selects the appropriate algorithm for each input fasta, and the latter is a recently-published method based on graph clustering.

We cleaned and trimmed alignments with GBLOCKS (v0.91b) (Castresana, 2000) and trimAl (v1.2) (Capella-Gutiérrez et al., 2009). For GBLOCKS we used the minimum (“-b2=0”) or default (85%) values for the minimum number of sequences for a flank position and allowed all gap positions (“-b5=a”). For trimAl we explored three different options: “gappyout”, “automated1” and “gapthreshold=0.2” following Portik & Wiens (2021).

Potentially paralogous sequences were flagged by HybPiper. Flagged sequences for each exon were collated, and RAxML (v8.2.9) (Stamatakis, 2014) was used to generate a phylogenetic tree with potentially paralogous sequences for each locus with flagged sequences. If there is true paralogy at a locus, flagged sequences would be expected to group by paralog, rather than by specimen. Phylogenetic trees were plotted and the presence of this pattern was examined by eye. Non-flagged sequences were not used to construct trees. If trees contained three or fewer specimens with flagged sequences, these loci were classified as non-paralogs.

On the other hand, if the locus contained large numbers of species with flagged sequences (i.e. >30) it was removed. If sequences were determined to be paralogous after examination, the entire locus was removed from the dataset. Note that paralogs could only be detected using those specimens that were run through the HybPiper pipeline (i.e., not including those where recoveries were previously done, such as 1KP data).

### Locus filtering

We performed two main filtering steps to minimize gene tree error and identify a reliable and high-quality set of loci for phylogenetic inference (Fig. 3). First, we removed sequences at each locus in which <10% of the length of the exon was recovered, before the alignment step. We also examined the effect this type of filtering had on phylogenetic reconstruction by varying the minimum percentage length threshold. The lowest value was 10% followed by 25% and 50% in the most extreme filtering.

**FIGURE 2.**
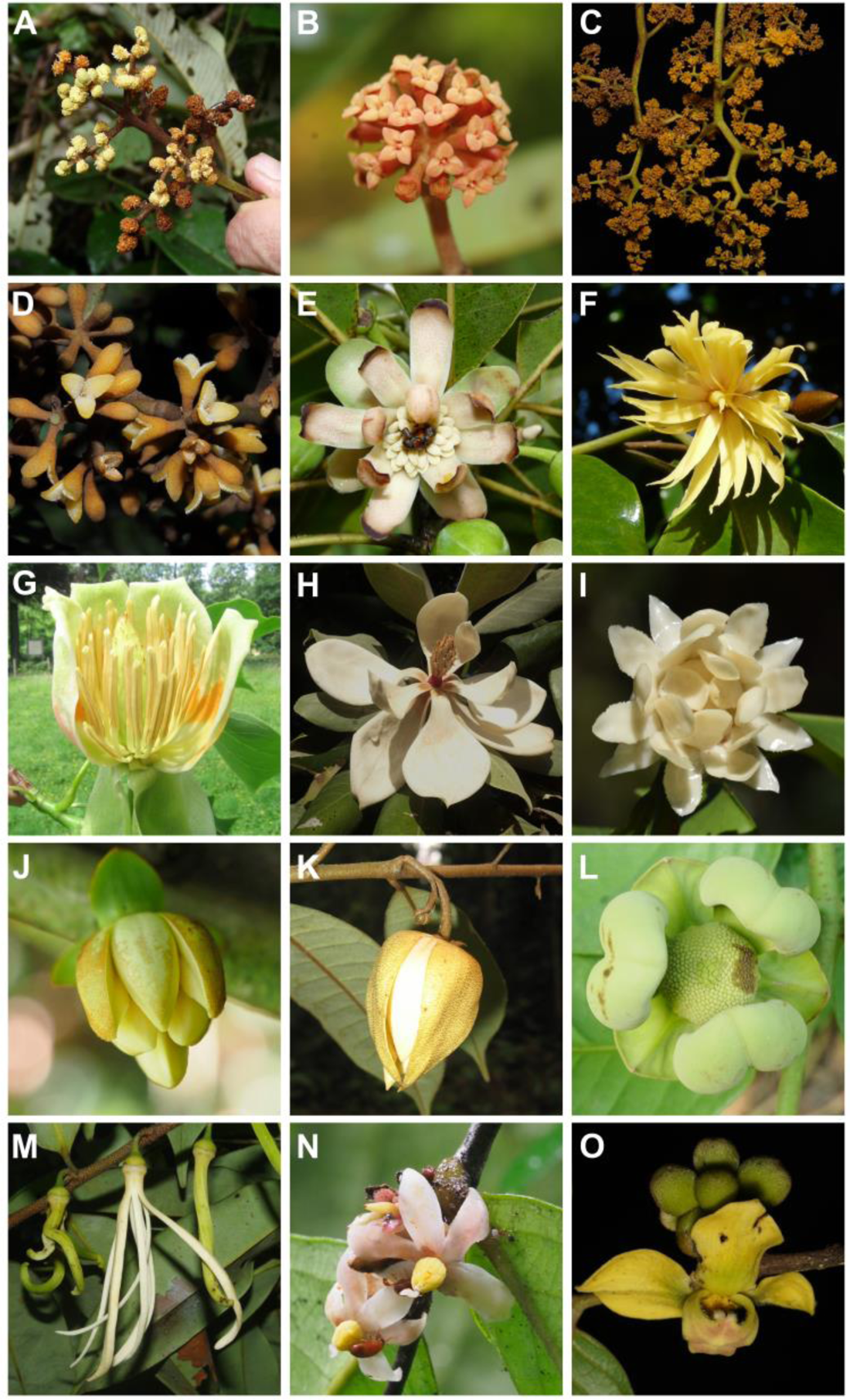
Floral diversity in Magnoliales. A) Myristicaceae, Mauloutchieae: *Pycnanthus angolensis* (Welw.) Warb.; B) Myristicaceae, Scyphocephalieae: *Scyphocephalium mannii* (Benth.) Warb.; C) Myristicaceae, Horsfieldieae: *Horsfieldia sylvestris* (Houtt.) Warb.; D) Myristicaceae, Viroleae: *Virola sebifera* Aubl.; E) Degeneriaceae: *Degeneria vitiensis* I.W.Bailey & A.C.Sm.; F) Himantandraceae: *Galbulimima baccata* F.M.Bailey; G) Magnoliaceae: *Liriodendron tulipifera* L.; H) Magnoliaceae: *Magnolia sororum* Seibert; I) Eupomatiaceae: *Eupomatia laurina* R.Br.; J) Annonaceae, Anaxagoreoideae: *Anaxagorea brevipes* Benth.; K) Annonaceae, Ambavioideae: *Meiocarpidium oliverianum* (Baill.) D.M.Johnson & N.A.Murray; L) Annonaceae, Annonoideae: *Cymbopetalum baillonii* R.E.Fr.; M) Annonaceae, Annonoideae: *Xylopia aethiopica* (Dunal) A.Rich.; N) Annonaceae, Malmeoideae: *Sirdavidia solannona* Couvreur & Sauquet; O) Annonaceae, Malmeoideae: *Mitrephora diversifolia* (Span.) Miq. Photo credits: A, E, F, I: Hervé Sauquet; B, D, G, J, K, L, M, N: Thomas L.P. Couvreur; C, H, O: Zacky Ezedin.

**FIGURE 3.**
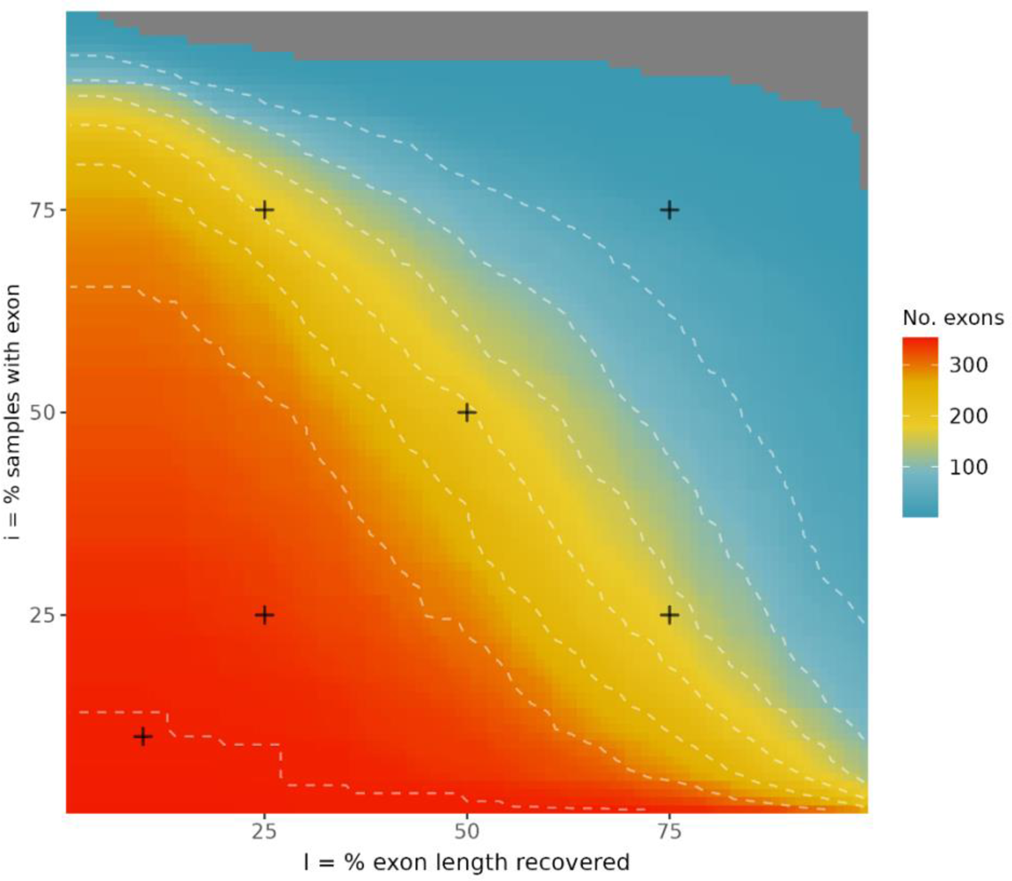
Heatmap of exon recovery. Each cell corresponds to the number of exons retained with different filtering thresholds. The X-axis is the minimum proportion of the exon length recovered (‘l’) and the Y-axis is the minimum number of samples this proportion was recovered in (‘i’). The dashed white lines are contours to show the slope of the surface. Crosses indicate the thresholds used to generate the range of trees assessed in this study.

Second, we chose a set of loci for phylogenetic inference depending on the proportion of specimens for which a given exon length threshold was achieved. For example, our main analyses are based on the set of loci for which at least 50% of the exon was recovered in at least 50% of specimens. Again, we varied these values from 10% to 25%, 50% and 75% for each to examine their effect on support values and tree topology. We also iterated through the two parameters to examine how they interact and how their variation changes the number of loci selected as reliable for phylogenetic inference. Again, all filtering steps only included those species for which the recoveries were done in this study.

### Phylogenetic inference

We constructed phylogenetic trees using both coalescent and concatenation approaches, though our assessment of different methodological approaches was based on coalescent approaches for computational feasibility and ease of interpretation. The coalescent approach was implemented in ASTRAL-III (v5.7.7) (Zhang et al., 2018), which takes gene trees as input to construct a species tree and is particularly useful for understanding gene tree discordance.

Gene trees were constructed for each locus using RAxML (v8.2.9) (Stamatakis, 2014) using the rapid bootstrap “-f a” option with 100 bootstrap replicates to provide support values for downstream filtering. For each tree we chose a GTRGAMMA model that uses empirical base frequencies, the gamma model of rate heterogeneity and the general time-reversible (GTR) matrix for substitution rates. Branches with bootstrap support <10% were collapsed using Newick Utilities program nw_ed (v1.6) (Junier and Zdobnov, 2010) to help improve the accuracy of downstream species tree inference. We then ran ASTRAL-III using the gene trees from our selected set of loci using options to output local posterior probability (LPP) “-t 3” and quartet support “-t 1”.

The concatenation approach was implemented in IQTREE (v2.1.2) (Minh et al., 2020). After removing potential paralogs and filtering the dataset, alignments were concatenated using the program phyx and its function ‘pxcat’ (Brown et al., 2017). A separate substitution model was applied to each locus to account for rate variation. We ran IQTREE with options “-m MFP+MERGE” to perform PartitionFinder (Lanfear et al., 2017) then tree reconstruction, and “-bb 1000” to perform 1000 ultrafast bootstraps.

We used DiscoVista (Sayyari et al., 2018) to examine how well species trees built with different sets of loci and alignment/trimming methods supported major clades in Magnoliidae. We then examined gene tree conflict in detail for two contentious relationships in our trees, the placement of Hydnoraceae in Piperales and of genus *Meiocarpidium* in Annonaceae. In addition, we calculated pairwise phylogenetic (Robinson-Foulds) distances among our different trees using the R package ‘treedist’ (Smith, 2022).

We used four different summary statistics to evaluate the quality of different approaches: (1) normalized scores output for each ASTRAL run that represent the percentage of gene trees that support the species tree; (2) the total number of quartets used in each analysis; (3) the mean Local Posterior Probability (LPP) across all nodes; and (4) the mean quartet support (QS), i.e. the proportion of quartets that support the node in the species tree, across all nodes. We also examined whether different approaches led to higher support at shallow vs deep nodes.

## RESULTS AND DISCUSSION

### Impact of filtering, locus choice, and methodology on phylogenetic reconstruction

We built a range of phylogenetic trees using 14-339 loci from the Angiosperms353 kit. The kit performed well across the different magnoliid orders, though we found little overlap between high-quality loci produced for Piperales compared to the other orders (Appendix S2). This may be in-part because fewer loci were recovered in Piperales (Table 1). A total of 11 putatively paralogous loci were detected and these were removed from all downstream analyses. In general, coalescent (using ASTRAL) and concatenation (using IQTREE) approaches yielded similar topologies (Fig. 5; Appendix S3, S4). The few notable differences concentrated in Aristolochiaceae (most notably the placement of Hydnoraceae), Piperales, Lauraceae, Myristicaceae, and various subclades of Annonaceae (Appendix S4). For the remainder of the results, we focus mostly on phylogenetic trees generated by ASTRAL as these allowed us to dive further into incongruence among different gene trees and difference in methodological approaches.

**FIGURE 4.**
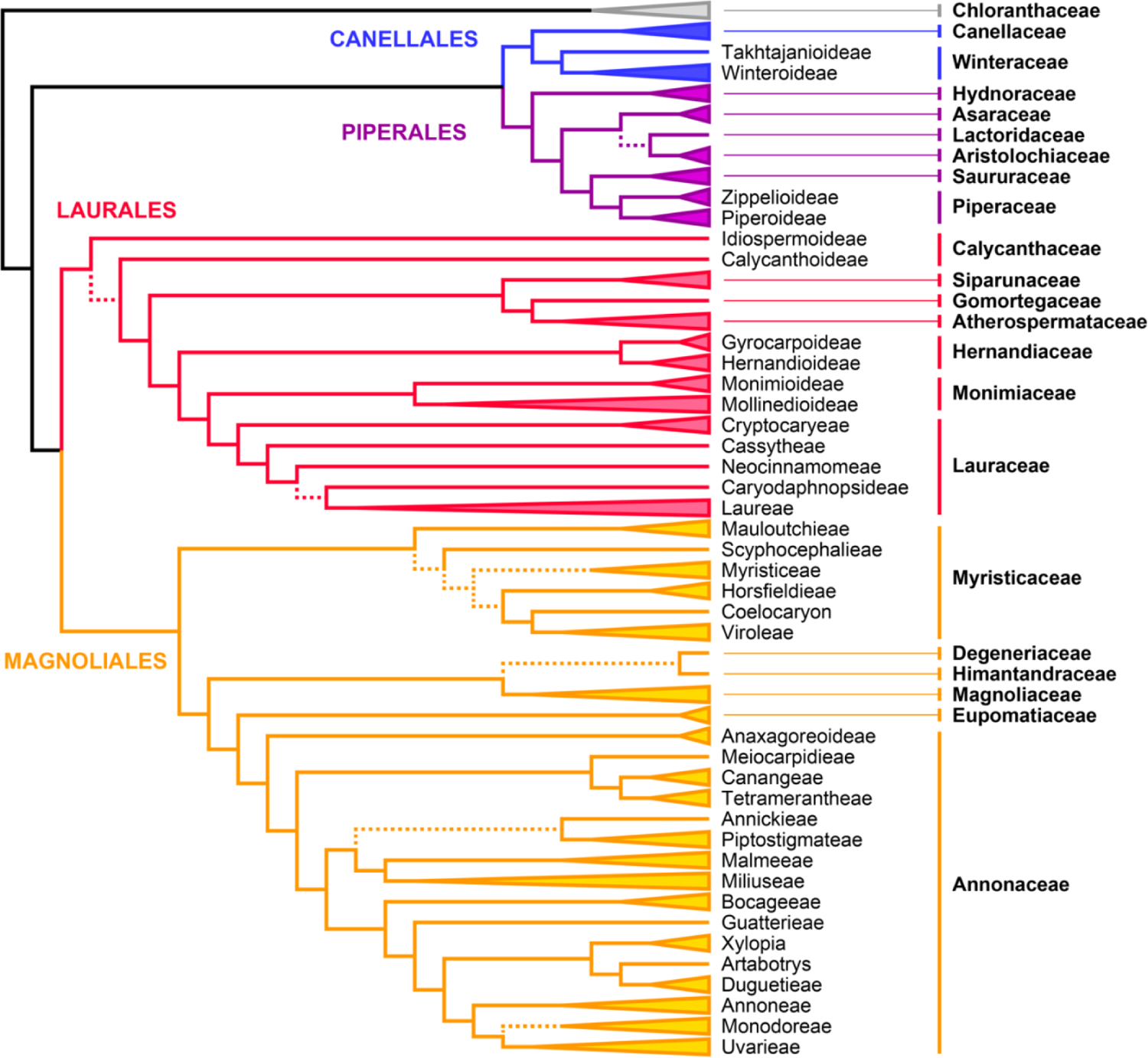
Summary of phylogenetic relationships in Magnoliidae obtained in the coalescent analysis of Angiosperms353 nuclear data. Triangle cartoons represent collapsed clades (for full detail, see Fig. 5). Tip and clade labels correspond to the updated phylogenetic classification of Magnoliidae presented in this paper. Dashed branches received local posterior probabilities <0.95 in this analysis.

**FIGURE 5.**
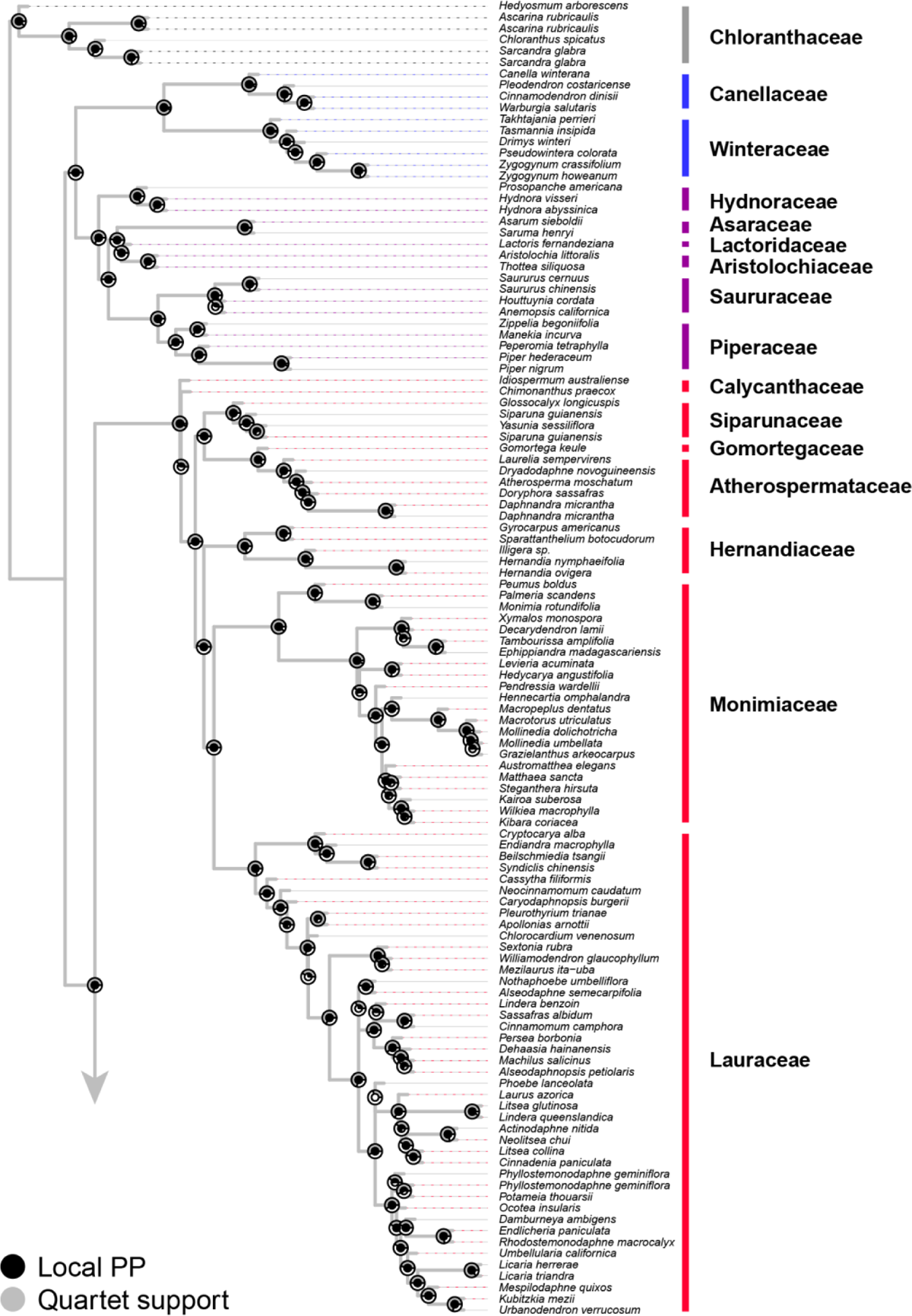
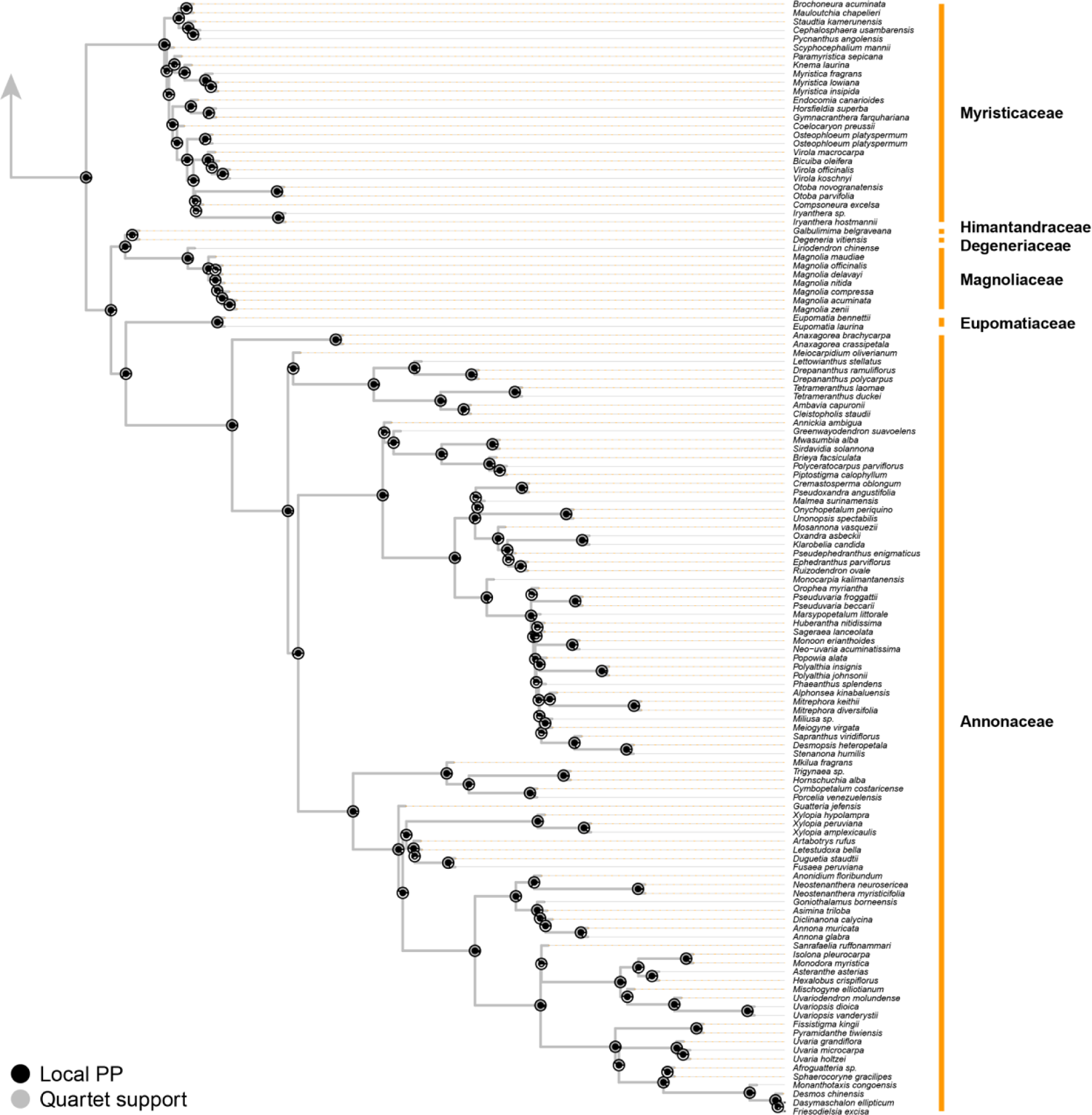
Phylogenetic relationships in Magnoliidae obtained in our main coalescent analysis (same tree as Fig. 4). This tree was reconstructed using the following methods: it was aligned with MAFFT, trimmed with gblocks option “–b2=0”, and the filtering thresholds of r = 10%, l = 50% and i = 50% were used. This tree achieved the highest combined quality score of those compared (Appendix S5). At each node the larger grey circle indicates quartet support (QS) and the inner black circle represents local posterior probability (LPP).

**TABLE 1.**
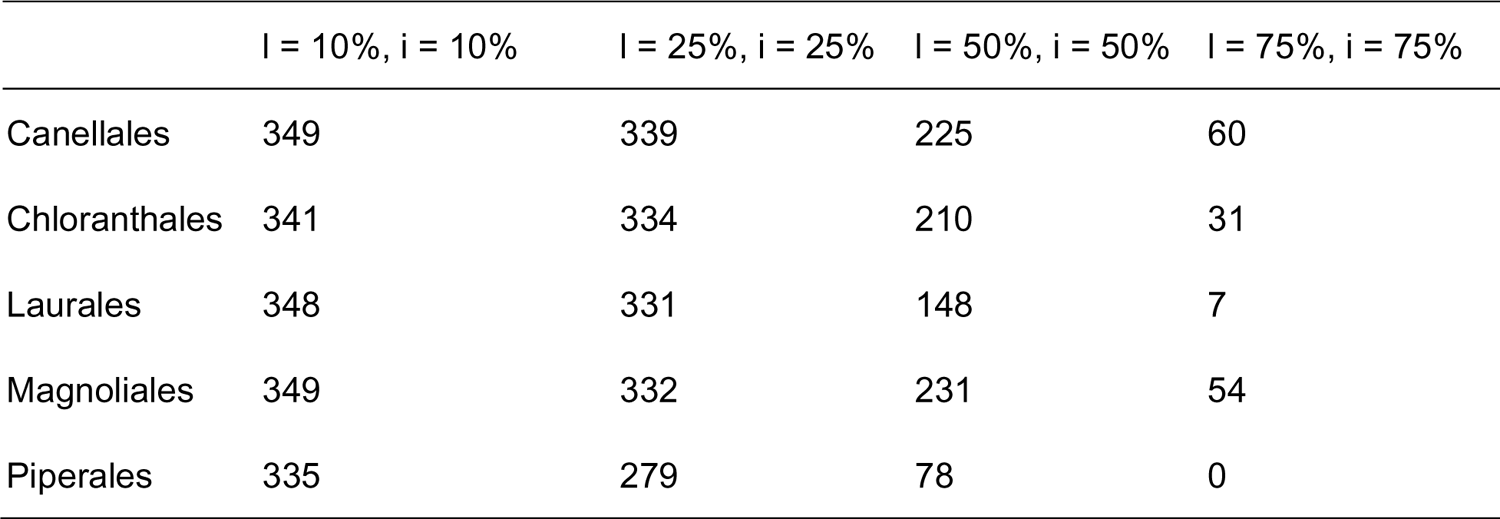
Numbers of exons selected as suitable for phylogenetic analyses per order. Each column indicates a different filtering threshold relating to the proportion of exon recovered (l) and the percentage of individuals achieving this proportion (i).

|  | l = 10%, i = 10% | l = 25%, i = 25% | l = 50%, i = 50% | l = 75%, i = 75% |
| --- | --- | --- | --- | --- |
| Canellales | 349 | 339 | 225 | 60 |
| Chloranthales | 341 | 334 | 210 | 31 |
| Laurales | 348 | 331 | 148 | 7 |
| Magnoliales | 349 | 332 | 231 | 54 |
| Piperales | 335 | 279 | 78 | 0 |

### Effects of filtering, locus choice and methodological choices on branch support

We examined the effect of changing how loci are filtered, sequences are aligned, and alignments are trimmed on tree topology and support. Loci were first filtered prior to alignment using a threshold ‘r’ to remove sequences with only short fragments of exons, from 10 to 50% of the reference length. As ‘r’ was increased this tended to increase mean QS and limit mean LPP (Appendix S5). This was likely because increasingly fewer quartets were available to evaluate species trees but those that remained were based on more data and tended to better support the species tree hypothesis.

Loci were then filtered based on whether a given proportion of the total length of exon was recovered (‘l’) in a given proportion of the total number of individuals (‘i’). We built a ‘locus surface’ to demonstrate how changing these two filtering criteria altered the number of loci retained (Fig. 3) and gives an indication of the overall quality of our data set. When both of the locus filtering criteria were high (l = 75% & i = 75%; the top right of Fig. 3) most of the loci were removed from the data set, leaving just 18 to build the tree (before paralog removal).

Decreasing the threshold on the proportion of individuals while maintaining large portions of the exons (l = 75% & i = 25%; bottom right of Fig. 3) substantially increased the number of loci to 168. Similarly, allowing for smaller portions of the exon to be used if they were present in the majority of individuals (l = 25% & i = 75%, top left of Fig. 3) led to 191 loci retained. Less stringent filtering thresholds for both criteria (l = 25% & i = 25% to l = 10% & i = 10%, bottom left of Fig. 3) led to the highest locus retention (338 and 350 respectively). The center of the space (l = 50% & i = 50%) yielded 203 loci. If strict filtering (i.e. l = 75% & i = 75%) was used there were not enough independent loci to have high confidence in the species tree topology, which resulted in a relatively low mean LPP and high QS, similar to when we increased ‘r’ (Appendix S5). Furthermore, this resulted in many nodes that were based on just a handful of loci, which can bias relationships if these happen to not match the true species tree. As filtering was relaxed and increasing numbers of loci were used the pattern reversed – the large number of loci used meant that LPP was high as confidence in the species tree topology increased but more conflict was also introduced, which lowered QS. Quartet scores were lowest (∼63%) in analyses where filtering was most relaxed (l = 25% & i = 25% or l = 10% & i = 10%; Appendix S5). We found that decreasing ‘l’ and ‘i’ seemed to have a greater effect on the number of loci kept in Piperales and Laurales when compared to the other orders (Table 1).

We used two aligners: MAFFT and MAGUS but these tended to have relative small effects on quality metrics (Appendix S5). Likewise, when only exons were used (instead of supercontigs) trees had marginally lower quality scores (LPP: 0.897 vs 0.914, QS: 63.041 vs 65.354). Modifying the approach to alignment trimming seemed to have a larger effect on the tree quality than other changes. Methods that were less conservative (i.e., a higher amount of less-well aligned data was kept) generally resulted in lower normalized scores but higher LPP, but this was only clear in extreme cases (e.g. trimAl -gapthreshold=0.2). Across all analyses, average LPP typically ranged from 0.88-0.92, QS varied from 63-65.5% and normalized scores from 0.912-0.961. This indicated that, generally, changing the methodological approach did not have a large effect on these support metrics. However, we found that the number of quartets used varied by more than 20-fold, from approximately 1.1e+9 to 22.2e+9 (Appendix S5). When examining the distribution of support within the trees, we found that uncertainty was generally concentrated in a few areas of the tree, regardless of the approach taken to build the tree. This was most evident in the Miliuseae (Annonaceae) and to a lesser extent in Mollinedioideae (Monimiaceae), Laureae (Lauraceae), and Myristicaceae (Fig. 5).These conflicts may be, at least in part, due to lower exon recovery in some of the species sampled in these clades (Appendix S6).

### Effects of filtering, locus choice and methodological choices on topology

Even when support values varied little, changes to methodology may have an effect on tree topology. We used DiscoVista to visualize how species trees built with different combinations of filtering criteria, aligners and trimmers outlined above affect the support for magnoliid families and orders (Appendix S7). Note that trees were evaluated after five taxa that tended to move among different analyses were removed (*Yasunia sessiliflora, Pleurothyrium trianae, Chlorocardium venenosum, Apollonias arnottii* and *Potameia thouarsii*). Most of these clades were strongly supported under the majority of scenarios. Exceptions included Aristolochiaceae, for which most analyses strongly rejected monophyly of the family, likely due to the placement of Hydnoraceae relative to the other species (Fig. 5, Appendix S7). Indeed, our IQTREE analysis actually provided strong support for the monophyly of Aristolochiaceae, with Hydnoraceae nested within (Appendix S3). There was also some uncertainty in Calycanthaceae as support was generally lower and rejected in a few cases (Appendix S7).

These results were generally mirrored when looking at the proportion of gene trees that supported major clades (Appendix S8) in our ASTRAL tree. Lauraceae had the highest number of gene trees that strongly rejected the monophyly of the family, though this was only roughly a quarter of the total number. For some small families (Siparunaceae and Calycanthaceae), the majority of loci could not be used to measure support as recoveries were poor. Similarly, few loci were available for Magnoliales likely because most loci were not present in the large number and highly divergent species required to validate this relationship.

To quantify how different analytical scenarios affect tree topology we calculated phylogenetic distance among our trees (Appendix S9). Distances appeared to be primarily dictated by ‘r’ and the alignment trimming algorithm. These two factors affect the amount of sequence data used to build a tree. In the most extreme cases they may lead to dropout if all sequences for an individual (or alignment) are removed from the data set, which likely contributes significantly to the calculation of tree distances. When we examined our trees more closely we found that differences in topology were consistently located in Myristicaceae, Miliuseae (Annonaceae) and Lauraceae, which were also generally the parts of the trees with low support. When ‘r’ was at its highest (r = 50%), major clades within Piperales were unstable or polyphyletic. When alignment trimmers and ‘r’ were changed, the placement of families Atherospermataceae, Hernandiaceae, and Siparunaceae also varied in some cases. Despite these differences we note that the majority of relationships remained stable despite methodological changes.

### Choosing the ‘best’ tree

The definition of what determines if one tree is better than another depends on the goals of the analysis. For example, one way would be to see which one provides the strongest support for the major clades in the group of interest (in this case these are magnoliid families and orders), in which case we may choose a tree based on exons as it had among the highest support across major clades (though others had similar levels of support; Appendix S7).

Exons may reconstruct these deeper relationships well as they do not contain the noise introduced by rapidly changing intronic regions, however they may be less useful for more recent divergences as they are generally highly conserved. Generally, analytical scenarios that were good at shallow node depths (<5) were also good at deeper node depths (≥5) but support was generally higher at shallow depths across all analyses (Appendix S10). As expected, the tree based on exon data alone had the lowest average QS across shallow nodes.

Alternatively, the metrics quantifying the level of support across all nodes of the tree can be used to assess their quality. However, this approach was made more complex because we found that different evaluation metrics were maximized with different analytical scenarios. The number of quartets was highest when filtering and trimming was less severe. This metric appeared to have weak, negative relationships (Appendix S11) with normalized scores (the proportion of gene tree quartets that match the species tree) and mean QS (i.e., these metrics were higher when fewer quartets were used). Scores and mean quartet support tended to be maximized when ‘l’ and ‘i’ filtering was most stringent. Mean LPP was positively correlated with normalized score and was maximized when trimming was done with trimAl ‘–gapthreshold=0.2’, which we highlighted above as a rather extreme approach when compared to the rest, likely removing a substantial amount of informative data. We note that high support does not guarantee that the relationship reflects the true species tree, particularly when only a handful of loci are available to reconstruct a relationship. However, given that the majority of nodes in most of our trees are supported by large numbers of rather large loci with independent histories, this type of bias is probably rare.

Overall, our results indicate that there was no single approach that was best, but instead several different ones were found that maximized different ways of assessing quality, albeit with relatively limited effect on support and topology. In an attempt to choose a single tree to represent relationships among magnoliid species we scaled and summed the four quality metrics used. We found that the tree presented in Figures 4 and 5 (mafft, gblocks option “–b2 0”, r10, i50, l50) had the highest total (Appendix S5), so we use this tree as the reference tree for the rest in this manuscript. However, we reiterate that that there remains some uncertainty about topology linked to methodological choices, as noted above. As a result, all remaining trees are provided as Appendix S12.

### Magnoliidae backbone

Our results lend further support to the split of Magnoliidae into two main clades: Canellales and Piperales, and Laurales and Magnoliales (Fig. 4). Furthermore, the four orders and all 18 families sensu APG IV (2016) were recovered as monophyletic across all our analyses, except for Aristolochiaceae, and Calycanthaceae in a few cases (see below; Fig. 5; Appendix S3, S7). These relationships emerged early in angiosperm molecular phylogenetics based on Sanger sequencing data and have remained stable ever since (Qiu et al., 1999; Sauquet et al., 2003; Soltis et al., 2011; Massoni et al., 2014). They were also confirmed by early and more recent phylogenomic studies based on plastid genomes (Moore et al., 2007; Li et al., 2019, 2021) or nuclear transcriptome data (Wickett et al., 2014; One Thousand Plant Transcriptomes Initiative, 2019). To the best of our knowledge, this study represents the first attempt to comprehensively sample all families of magnoliids (and most genera) in a phylogenomic framework based on nuclear genome data.

Within orders, our results broadly corroborate the relationships among families identified by previous studies, but also suggest potential solutions to long-term conflicts. In Piperales, our data suggest a new placement for the holoparasitic Hydnoraceae, with implications for the phylogenetic classification of the order. Indeed, a key question remains regarding the relationships among the traditional Aristolochiaceae, Lactoridaceae, and Hydnoraceae sensu APG III and previous classifications (APG III, 2009). Previous work had suggested that both Lactoridaceae and Hydnoraceae are nested in Aristolochiaceae (Naumann et al., 2013; Massoni et al., 2014), leading to the merging of the three families into a broader Aristolochiaceae (APG IV, 2016). However, a recent study based on data from all three plant genomes proposed instead to recognize four families (Aristolochiaceae, Asaraceae, Lactoridaceae, and Hydnoraceae) to account for both the high uncertainty persisting regarding their relationships and the strong morphological differentiation among these four clades (Fig. 1C-G; Jost et al., 2021). Our coalescent analyses suggest that Hydnoraceae may be the sister group to all remaining Piperales. The most frequent topology (50%) in the gene trees used to build our main tree (Fig. 5) supported Hydnoraceae (*Hydnora* and *Prosopanche*) being distinct from other Aristolochiaceae sensu APG IV (Fig. 6a). In contrast, our concatenation analysis (and a few coalescent ones) suggests a position nested in Aristolochiaceae sensu APG IV (Appendix S3), consistent with the main (137-loci) analysis of Jost et al. (2021). This was also the second most frequent (35%) topology in our main coalescent analysis (Fig. 6a), indicating that substantial conflict exists, even if the species tree topology is well-supported.

**FIGURE 6.**
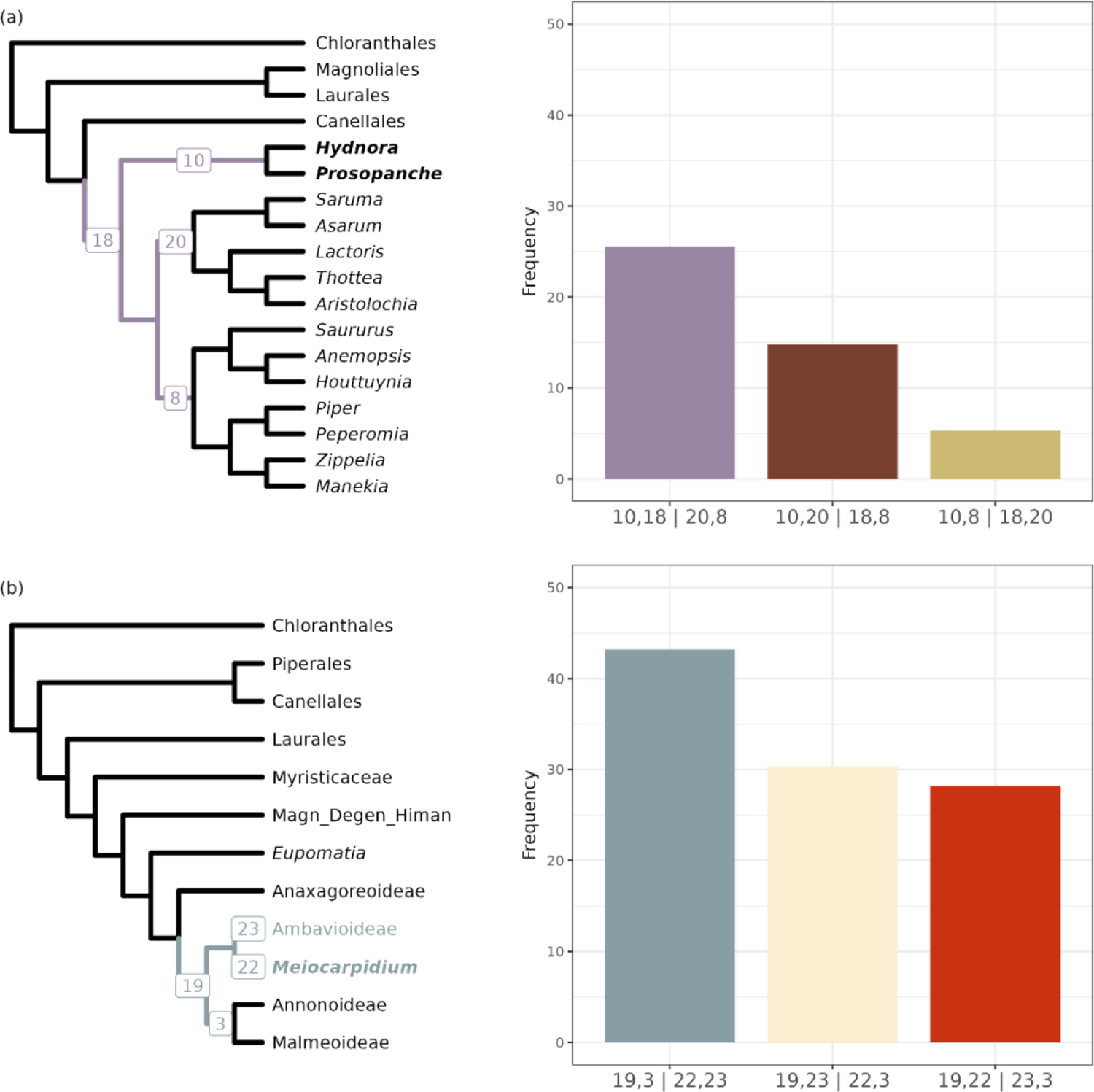
Topology test to understand gene tree conflict for the placement of (a) Hydnoraceae and (b) *Meiocarpidium*. The left side is a summary tree where the relevant branches are colored and the focal taxa in bold. Branch labels correspond to those used in the X axis labels in the barplots on the right. The barplots show the frequency of each quartet, labelled below the bar. For the tree in (b) we combined Magnoliaceae, Himantandraceae and Degeneriaceae into a single tip.

In Laurales, our results highlight conflicting relationships among Hernandiaceae, Lauraceae, and Monimiaceae, consistent with previous studies which have found it difficult to resolve this question (Doyle and Endress, 2000; Renner and Chanderbali, 2000; Massoni et al., 2014). Both our coalescent and concatenation analyses suggest Hernandiaceae to be sister to a clade comprising Lauraceae and Monimiaceae, unlike the majority of previous studies, in which Lauraceae were found to be sister to a clade including Hernandiaceae and Monimiaceae (Qiu et al., 1999; Soltis et al., 2011; Li et al., 2019, 2021; One Thousand Plant Transcriptomes Initiative, 2019). Likewise, analyses of morphological data strongly favored Monimiaceae to be sister to a clade of Hernandiaceae and Lauraceae (Doyle and Endress, 2000, 2010; Sauquet et al., 2003). However, despite the apparent support for this relationship in the analyses presented here, these results should be taken with caution because it remained unstable across different analytical approaches.

In Magnoliales, our data provide new support for the placement of Magnoliaceae as sister to a clade of Degeneriaceae and Himantandraceae (Fig. 5; Appendix S3). Previous studies that sampled the three families had either suggested the same arrangement with low support (reviewed by Massoni et al., 2014) or left these relationships unresolved (Qiu et al., 1999; Soltis et al., 2011; Massoni et al., 2014; Li et al., 2021), while combined analyses of molecular and morphological data had supported instead Magnoliaceae to be sister to a broader clade of Degeneriaceae, Himantandraceae, Eupomatiaceae, and Annonaceae (Doyle and Endress, 2000; Sauquet et al., 2003).

### Relationships within families

#### Monimiaceae

The phylogenetic tree recovered two subfamilies for Monimiaceae, Monimioideae and Mollinedioideae, both with high support (Fig. 5; Appendix S3). Renner et al. (2010) also recovered the same topology based on two cpDNA and one rDNA regions, as did Massoni et al. (2014) with six cpDNA regions, four mDNA, and two rDNA regions. *Hortonia*, the only genus of subfamily Hortonioideae, was initially sampled but resulted in low quality data and had to be excluded from our analyses. The genus was sampled in previous studies and was recovered as sister to Mollinedioideae with high support (Renner et al., 2010; Massoni et al., 2014). Relationships among the three genera of Monimioideae are consistent with previous work (Renner et al., 2010; Massoni et al., 2014), with *Peumus* sister to a clade comprising *Monimia* and *Palmeria*.

In Mollinedioideae, tribe Hedycaryeae was recovered as paraphyletic with *Xymalos, Decarydendron, Tambourissa*, and *Ephippiandra* as sister to a clade including *Hedycarya* and *Levieria*, and other Mollinedioideae genera, however, with low support and unstable across our analyses (Fig. 5; Appendix S3). Massoni et al. (2014) also recovered *Xymalos* in a clade with *Decarydendron* and *Tambourissa*. Instead, Renner et al. (2010) had found *Xymalos* as sister to all other Mollinedioideae genera. The presence of persistent tepals and fruiting receptacle not enclosing the drupelets support our findings positioning this genus as closely related to *Decarydendron* and *Tambourissa*.

The clade sister group to *Hedycarya* and *Levieria* includes all the genera assigned by Philipson (1987, 1988, 1993) to tribes Mollinedieae and Hennecartieae (with only the genus *Hennecartia*). Mollinedieae was recovered as paraphyletic due the presence of Hennecartieae as the sister group of the Neotropical Mollinedioideae clade. The circumscription of Hennecartieae relied on its discoid staminate flowers, anthers dehiscent by an equatorial slit, and pistillate flowers urceolate with oblate tepals (Philipson, 1988, 1993). Despite its peculiar morphology, *Hennecartia* was also recovered as sister to the other Neotropical Mollinedieae genera by Renner et al. (2010), but it appeared as a sister group of *Wilkiea* from Australia and Southeast Asia in Massoni et al. (2014). The enigmatic genera *Grazielanthus* and *Macrotorus* were placed in a clade with the remaining Neotropical Mollinedieae genera, as in Renner et al. (2010). Mollinedieae was circumscribed by anthers with longitudinal slits, pistillate receptacle opening by abscission of the upper part, and fruits exposed and attached to a reflexed fruiting receptacle enlarged. The genus *Macrotorus* was considered by Philipson (1987, 1988, 1993) as not sufficiently known to be placed because, at the moment, only staminate flowers were known. The placement of *Macrotorus* in tribe Mollinedieae was accepted by subsequent authors (Peixoto and Pereira-Moura, 2008; Lírio et al., 2015, 2020) based on its flower and fruit features. The monotypic genus *Grazielanthus* had not yet been described when Philipson (1987, 1988, 1993) revised the classification of Monimiaceae, however it was considered by Peixoto & Pereira-Moura in 2008 as a member of Mollinedieae despite its non-calyptrate flowers, because all the other characteristics of the genus supported this placement.

#### Lauraceae

Our analyses recovered the main tribes of Lauraceae recognized in previous work. Hypodaphnideae, represented by the monotypic genus *Hypodaphnis*, was not included in our analyses due to low quality data recovered. Cryptocaryeae (including *Cryptocarya, Endiandra, Syndiclis*, and *Beilschmiedia*) was recovered with high support, as in previous studies (Rohwer, 2000; Chanderbali et al., 2001; Li et al., 2020; Song et al., 2020). This tribe appears as the sister group to all other Lauraceae genera. Three monogeneric tribes were recovered as successive sister lineages to the remaining Lauraceae genera: Cassytheae (*Cassytha*), Neocinnamomeae (*Neocinnamomum*), and Caryodaphnopsideae (*Caryodaphnopsis*). These results generally agree with previous work (Rohwer, 2000; Chanderbali et al., 2001; Song et al., 2020), except for the relative positions of Cassytheae and Neocinnamomeae obtained by Chanderbali et al. (2001).

Tribe Cinnamomeae (as circumscribed by Song et al. 2020) was recovered as monophyletic, but with low support (Fig. 5; Appendix S3). The genera *Sextonia, Williamodendron*, and *Mezilaurus* form a clade sister to a clade comprising the *Persea-Cinnamomum* group and other Cinnamomeae genera. The *Persea*-*Cinnamomum* lineage is sister to a clade including *Laurus, Neolitsea,* and allies, and the clade comprising *Damburneya, Licaria,* and allies. The genera *Chlorocardium, Phoebe,* and *Yasunia* appeared unstable in our trees.

#### Myristicaceae

Phylogenetic relationships within Myristicaceae allow for the first time the delimitation of five tribes based on clades recovered with high support. A strongly supported clade of five Malagasy and African genera (i.e., *Brochoneura, Cephalosphaera, Mauloutchia, Pycnanthus*, and *Staudtia*) is found to be sister to the remainder of the family. All of these genera plus genus *Doyleanthus* (not sampled here) were previously recovered in a monophyletic group informally named the ‘mauloutchioids’ (Sauquet et al., 2003; Massoni et al., 2014; Frost et al., 2022). This clade is formally described here as tribe Mauloutchieae (Appendix 1), including *Doyleanthus* based on previous work (Sauquet et al., 2003). Within this tribe, the Malagasy (*Brochoneura, Mauloutchia*) and African (*Cephalosphaera, Pycnanthus, Staudtia*) genera form two strongly supported clades, whereas the latter was previously recovered as a grade (Sauquet et al., 2003). Of all the tribes, Mauloutchieae exhibits by far the widest range of morphological diversity, particularly in anther types, fruit ornamentation, dehiscence, aril type, and endosperm rumination (Sauquet et al., 2003).

The Central African *Scyphocephalium* appears isolated as sister to the remaining of the family in our coalescent tree (Fig. 5). Although the genus is resolved in a different position in our concatenation analysis (Appendix S3), namely nested in the Mauloutchieae together with *Paramyristica*, we treat this alternative position, for both taxa, as a likely reconstruction artifact. Due to its topological isolation along with its unique inflorescence morphology (Fig. 2B), *Scyphocephalium* is here classified into a separate tribe, Scyphocephalieae.

The Asian genera were recovered in two well supported clades, but with uncertain relationships. For this reason, the two Asian clades are here recognized as separate tribes: Horsfieldieae and Myristiceae (Appendix 1). Generic relationships within Horsfieldieae are well supported as [*Endocomia* [*Gymnacranthera* + *Horsfieldia*]]. Myristiceae is represented by the sister pair of *Knema* and *Myristica*, a relationship also recovered in earlier analysis of molecular data (Sauquet et al., 2003), along with the monotypic genus *Paramyristica* in our coalescent tree (Fig. 5). Despite the alternative placement of this genus in the concatenation analysis (Fig. S3), we tentatively assign *Paramyristica* to Myristiceae.

*Coelocaryon* is isolated in all of our analyses and consistently appears as sister to Viroleae. However, its position is poorly supported and appears erratic. For example, when exon filtering in coalescent analyses is changed to 75%, it appears sister to Myristiceae with poor support. The well-supported sister relationship between *Coelocaryon* and *Pycnanthus* seen in prior plastid analyses (Sauquet et al., 2003; Frost et al., 2022) was not recovered. Due to this conflict, we believe further data will be required before assigning *Coelocaryon* to any tribe.

All the Neotropical genera formed a well-supported clade, here recognized as tribe Viroleae. Within this tribe, relationships remain largely unclear with no single topology found to be well supported. The Brazilian endemic genus *Bicuiba*, originally described as a species of *Virola* then subsequently segregated out as a monotypic genus by de Wilde (1991) on the basis of inflorescence structure, appears to be nested in *Virola*. A recently published molecular study on *Otoba* resolved this genus as sister to the rest of the tribe, albeit with poor support (posterior probability <50) (Frost et al., 2022). In previous phylogenetic studies, *Otoba* was recovered in a clade with the African genera *Coelocaryon* and *Pycnanthus* (Sauquet et al., 2003; Massoni et al., 2014).

*Haematodendron*, a monotypic Malagasy genus, was not included in this study and its placement is not yet resolved. Previously, it was recovered as sister to either *Bicuiba* (Sauquet et al., 2003) or *Staudtia* (Massoni et al., 2014), in both instances with poor support.

*Haematodendron* and other Malagasy genera of Mauloutchieae share similarities of the androecia, pollen, and aril, but differ in floral shape and endosperm ruminations (Kühn and Kubitzki, 1993). Therefore, until further information becomes available, *Haematodendron* remains incertae sedis at tribal level.

#### Annonaceae

The inferred phylogenetic relationships within Annonaceae are broadly similar to those inferred using an Annonaceae specific nuclear probe set (Couvreur et al., 2019) and plastid data (Chatrou et al., 2012; Guo et al., 2017). Interestingly, there is little overlap in terms of regions between the two probe kits, with only 12 Annonaceae loci clustering with regions of the Angiosperms353 kit. This would also suggest that the probe set of Couvreur et al. (2019) could be combined with the Angiosperms353 kit (Johnson et al., 2019) to increase resolution in certain tribes such as Miliuseae and Malmeeae. Such an approach has already been undertaken in families such as Brassicaceae (Hendriks et al., 2021).

We retrieved all four previously described subfamilies as monophyletic and well-supported, namely Anaxagoreoideae, Ambavioideae, Annonoideae and Malmeoideae. Phylogenetic relationships among genera are largely consistent with previously published results (Couvreur et al., 2019; Chatrou et al., 2012; Guo et al., 2017) and therefore we will address only cases which diverge from previous findings.

The central African monotypic genus *Meiocarpidium* was recovered as sister to the rest of Ambavioideae with good support (1.0 LPP). However, we found substantial gene tree conflict in this relationship (Fig. 6b). The most frequent quartet topology (45%) was only slightly more common than the others (36%) but this was underlined by a large number of loci, resulting in high confidence in the species tree topology. *Meiocarpidium* was included in Ambavioideae in all but one of our filtering scenarios. Under the most stringent scenario (astral_r10_l75_i75_auto_b2_0), *Meiocarpidium* was recovered as sister to Annonoideae and Malmeoideae, however, as indicated above this filter setting result in very few markers retained.

The position of *Meiocarpidium* has been debated over the years with some authors suggesting it should be assigned to its own subfamily, Meiocarpidioideae (Chaowasku, 2020). This decision was based on a phylogenetic tree obtained from the analysis of eight plastid markers, which had inferred *Meiocarpidium* as sister to the rest of Annonaceae excluding *Anaxagorea* with mixed support values (Chaowasku, 2020), thus rendering Ambavioideae paraphyletic. This result was, however, never inferred in previous phylogenetic studies of the family also based on plastid markers. *Meiocarpidium* was always inferred as sister to the rest of the Ambavioideae with different levels of support from weak to strong (Couvreur et al., 2008, 2019; Chatrou et al., 2012; Guo et al., 2017). Given the results presented here and those based independently on the Annonaceae specific probes (Couvreur et al., 2019), we do not advocate for the establishment of subfamily Meiocarpidioideae. Furthermore, adopting a subfamily view would not be consistent with other similar cases across the family. For instance, the genus *Annickia* was inferred as sister to the rest of Malmeoideae, but was recognized at tribe level (Annickieae), and not subfamily level (Couvreur et al., 2019). Thus, we propose that the monotypic genus *Meiocarpidium* is instead assigned to the new tribe Meiocarpidieae within Ambavioideae, in addition to the two that already exist (i.e., Tetramerantheae and Canangeae; see Appendix 1).

One surprising outcome of our nuclear phylogenetic analysis (Fig. 5) compared to previous plastid phylogenetic studies is the non-monophyly of tribe Xylopieae (Chatrou et al., 2012; Couvreur et al., 2019). This tribe is composed of two genera, *Xylopia* a pantropical genus of trees, and *Artabotrys* a paleotropical genus of lianas. Plastid phylogenetic trees of Annonaceae have consistently inferred a strongly supported sister relationship between the two genera, both in family-level (one species per genus sampled) (Chatrou et al., 2012; Guo et al., 2017) and tribal-level (several species per genus) studies (Thomas et al., 2015). This result could be confirmed by increasing species sampling of *Artabotrys* and tribe Duguetieae using the Angiosperms353 and/or Annonaceae probe sets.

Finally, *Sanrafaelia* was inferred as sister to tribe Uvarieae, confirming the results of Couvreur et al. (2019), but in contrast with the results based on plastid markers (Couvreur et al., 2008, 2019; Chatrou et al., 2012; Guo et al., 2017). *Sanrafaelia* and its sister genus *Ophrypetalum* (not sampled here) were recently transferred to a new tribe based on a phylogenetic analysis of tribe Monodoreae using the Annonaceae probe kit (Dagallier et al., 2023).

## MAGNOLIIDAE PHYLOGENETIC CLASSIFICATION

Here we propose an updated phylogenetic classification for Magnoliidae consistent with the nuclear phylogenomic analyses presented in this paper (Fig. 3) as well as recent phylogenetic studies conducted with a variety of sequencing approaches in the larger families. In doing so, we aim at maximum stability by following recent phylogenetic classifications presented at the angiosperm level by the Angiosperm Phylogeny Group and at the family level by experts of these clades.

Briefly, we follow APG IV (2016) for orders and families, except for Aristolochiaceae for which we recognize four separate families following Jost et al. (2021). For Piperaceae, we follow Samain et al. (2008). For Hernandiaceae, we follow Michalak et al. (2010) in recognizing *Hazomalania* as a distinct genus. For Monimiaceae, we recognize the same three subfamilies as Philipson (1987, 1988, 1993), but none of the tribes at this stage as doing so would require additional sampling and resolution among genera of Mollinedioideae. For Lauraceae, we follow strictly Song et al. (2020) in recognizing six tribes. We acknowledge that, given the size of the family, it would seem more appropriate to recognize subclades at the subfamily level instead.

However, we feel it would be premature to propose an alternative to the latest phylogenetic system published by experts of the family, given our limited sample of genera here. For Myristicaceae, we propose the first formal infrafamilial classification by describing five new tribes, based in part on the informal clade names introduced by Sauquet et al. (2003). Lastly, for Annonaceae we follow the well-established system of subfamilies and tribes of Chatrou et al. (2012), with the addition of four new tribes introduced in more recent studies (Guo et al., 2017; Couvreur et al., 2019; Chaowasku, 2020) and Meiocarpidieae recognized here for the first time. The taxonomic treatment for the new tribes in Myristicaceae and Annonaceae is provided in Appendix 1. Taxa not sampled in this study are marked with an asterisk.

### Order Canellales Cronquist Family Canellaceae Martius

Canella, Cinnamodendron, Cinnamosma*, Pleodendron, Warburgia

### Family Winteraceae Lindley

Subfamily Takhtajanioideae Leroy

#### Takhtajania

Subfamily Winteroideae Arnott

Drimys, Pseudowintera, Tasmannia, Zygogynum

### Order Piperales Dumortier Family Hydnoraceae C.Agardh

Hydnora, Prosopanche

### Family Asaraceae Ventenat

Asarum, Saruma

### Family Lactoridaceae Engler

Lactoris

### Family Aristolochiaceae Jussieu

Aristolochia, Thottea

### Family Saururaceae Richard

Anemopsis, Gymnotheca*, Houttuynia, Saururus

### Family Piperaceae Giseke

Subfamily Verhuellioideae* Samain & Wanke

*Verhuellia\**

Subfamily Zippelioideae Samain & Wanke

Manekia, Zippelia

Subfamily Piperoideae Arnott

Peperomia, Piper

### Order Laurales Berchtold & Presl Family Calycanthaceae Lindley

Subfamily Calycanthoideae Burnett

Calycanthus, Chimonanthus

Subfamily Idiospermoideae Thorne

Idiospermum

### Family Siparunaceae (A.DC.) Schodde

Glossocalyx, Siparuna

### Family Gomortegaceae Reiche

Gomortega

### Family Atherospermataceae R.Brown

*Atherosperma, Daphnandra, Doryphora, Dryadodaphne, Laurelia, Laureliopsis*, Nemuaron\**

### Family Hernandiaceae Blume

Subfamily Gyrocarpoideae Pax

Gyrocarpus, Sparattanthelium

Subfamily Hernandioideae

Hazomalania*, Hernandia, Illigera

### Family Monimiaceae Jussieu

Subfamily Monimioideae Rafinesque

Monimia, Palmeria, Peumus

Subfamily Hortonioideae* Thorne & Reveal

*Hortonia\**

Subfamily Mollinedioideae Thorne

Austromatthaea, Decarydendron, Ephippiandra, Grazielanthus, Hedycarya, Hemmantia*, Hennecartia, Kairoa, Kibara, Kibaropsis*, Lauterbachia, Levieria, Macropeplus, Macrotorus, Matthaea, Mollinedia, Parakibara*, Pendressia, Steganthera, Tambourissa, Wilkiea, Xymalos

### Family Lauraceae Jussieu

Tribe Hypodaphnideae* Kosterm. Ex Reveal

*Hypodaphnis\**

Tribe Cryptocaryeae Nees

Aspidostemon*, Beilschmiedia, Cryptocarya, Dahlgrenodendron*, Endiandra*, Eusideroxylon*, Potameia*, Potoxylon*, Ravensara*, Sinopora*, Syndiclis, Triadodaphne*, Yasunia

Tribe Cassytheae Dumortier

Cassytha

Tribe Neocinnamomeae Yu Song, W.B.Yu & Y.H.Tan

Neocinnamomum

Tribe Caryodaphnopsideae Yu Song, W.B.Yu & Y.H.Tan

Caryodaphnopsis

Tribe Laureae Maout & Decaisne

Actinodaphne, Adenodaphne*, Aiouea, Alseodaphne, Alseodaphnopsis, Anaueria*, Aniba*, Apollonias, Chlorocardium, Cinnadenia, Cinnamomum, Clinostemon*, Damburneya, Dehaasia, Dicypellium*, Dodecadenia*, Endlicheria, Iteadaphne*, Kubitzkia, Laurus, Licaria, Lindera, Litsea, Machilus, Mespilodaphne, Mezilaurus, Mocinnodaphne*, Nectandra*, Neolitsea, Nothaphoebe, Ocotea*, Paraia*, Parasassafras*, Persea, Phoebe, Phyllostemonodaphne, Pleurothyrium, Povedadaphne*, Rhodostemonodaphne, Sassafras, Sextonia, Sinosassafras*, Umbellularia, Urbanodendron, Williamodendron

### Order Magnoliales Bromhead Family Myristicaceae R.Brown

Tribe Mauloutchieae Ezedin & Sauquet, trib. nov.

Brochoneura, Cephalosphaera, Doyleanthus*, Mauloutchia, Pycnanthus, Staudtia

Tribe Scyphocephalieae Ezedin & Sauquet, trib. nov.

Scyphocephalium

Tribe Horsfieldieae Ezedin & Sauquet, trib. nov.

Endocomia, Gymnacranthera, Horsfieldia

Tribe Myristiceae Ezedin & Sauquet, trib. nov.

Knema, Myristica, Paramyristica

Tribe Viroleae S.M.Oliveira & Sauquet, trib. nov.

Bicuiba, Compsoneura, Iryanthera, Osteophloeum, Otoba, Virola

Incertae sedis

*Coelocaryon, Haematodendron\**

### Family Degeneriaceae I.W.Bailey & A.C.Smith

Degeneria

### Family Himantandraceae Diels

Galbulimima

### Family Magnoliaceae Jussieu

Liriodendron, Magnolia

### Family Eupomatiaceae Orban

Eupomatia

### Family Annonaceae Jussieu

Subfamily Anaxagoreoideae Chatrou, Pirie, Erkens & Couvreur

Anaxagorea

Subfamily Ambavioideae Chatrou, Pirie, Erkens & Couvreur Tribe Meiocarpidieae Chatrou, Couvreur & Erkens, trib. nov.

Meiocarpidium

Tribe Canangeae Chaowasku

Cananga*, Cyathocalyx*, Drepananthus, Lettowianthus

Tribe Tetramerantheae R.E.Fr. ex Reveal

Ambavia, Cleistopholis, Mezzettia*, Tetrameranthus

Subfamily Annonoideae Raf. Tribe Bocageeae Endl.

Bocagea*, Cardiopetalum*, Cymbopetalum, Froesiodendron*, Hornschuchia, Mkilua, Porcelia, Trigynaea

Tribe Guatterieae Hook.f. & Thomson

Guatteria

Tribe Xylopieae Endl.

Artabotrys, Xylopia

Tribe Duguetieae Chatrou & R.M.K.Saunders

*Duckeanthus*, Duguetia, Fusaea, Letestudoxa, Pseudartabotrys\**

Tribe Annoneae Endl.

Annona, Anonidium, Asimina, Diclinanona, Disepalum*, Goniothalamus, Neostenanthera

Tribe Monodoreae Baill.

Asteranthe, Dennettia*, Hexalobus, Isolona, Lukea*, Mischogyne, Monocyclanthus*, Monodora, Ophrypetalum*, Sanrafaelia, Uvariastrum*, Uvariodendron, Uvariopsis

Tribe Uvarieae Hook.f. & Thomson

Afroguatteria, Cleistochlamys*, Dasymaschalon, Desmos, Dielsiothamnus*, Fissistigma, Friesodielsia, Monanthotaxis, Pyramidanthe, Schefferomitra*, Sphaerocoryne, Toussaintia*, Uvaria

Subfamily Malmeoideae Chatrou, Pirie, Erkens & Couvreur Tribe Annickieae Couvreur

Annickia

Tribe Piptostigmateae Chatrou & R.M.K.Saunders

Brieya, Greenwayodendron, Mwasumbia, Piptostigma, Polyceratocarpus, Sirdavidia

Tribe Malmeeae Chatrou & R.M.K.Saunders

Bocageopsis*, Cremastosperma, Ephedranthus, Klarobelia, Malmea, Mosannona, Onychopetalum, Oxandra, Pseudephedranthus, Pseudomalmea*, Pseudoxandra, Ruizodendron, Unonopsis

Tribe Maasieae* Chatrou & R.M.K.Saunders

*Maasia\**

Tribe Fenerivieae* Chatrou & R.M.K.Saunders

*Fenerivia\**

Tribe Phoenicantheae* X. Guo & R.M.K.Saunders

*Phoenicanthus\**

Tribe Dendrokingstonieae* Chatrou & R.M.K.Saunders

*Dendrokingstonia\**

Tribe Monocarpieae Chatrou & R.M.K.Saunders

Leoheo*, Monocarpia

Tribe Miliuseae Hook.f. & Thomson

*Alphonsea, Desmopsis, Huberantha, Marsypopetalum, Meiogyne, Miliusa, Mitrephora, Monoon, Neo-uvaria, Orophea, Phaeanthus, Platymitra*, Polyalthia, Polyalthiopsis*, Popowia, Pseuduvaria, Sageraea, Sapranthus, Stelechocarpus*, Stenanona, Tridimeris*, Trivalvaria*, Wangia*, Winitia*, Wuodendron\**

## ACKNOWLEDGMENTS

This work was funded by grants from the Calleva Foundation to the Plant and Fungal Trees of Life (PAFTOL) project at Royal Botanic Gardens, Kew. We would like to acknowledge the contribution of the Genomics for Australian Plants Framework Initiative consortium (https://www.genomicsforaustralianplants.com/consortium/) in the generation of data used in this publication. The Initiative is supported by funding from Bioplatforms Australia (enabled by NCRIS), the Ian Potter Foundation, Royal Botanic Gardens Foundation (Victoria), Royal Botanic Gardens Victoria, the Royal Botanic Gardens and Domain Trust, the Council of Heads of Australasian Herbaria, CSIRO, Centre for Australian National Biodiversity Research and the Department of Biodiversity, Conservation and Attractions, Western Australia.

## AUTHOR CONTRIBUTIONS

All authors jointly designed this study. Z.E., E.J.L., S.M.O., L.W.C., R.H.J.E., I.L., K.V., O.M., S.R., A.R.Z., W.J.B., T.L.P.C., F.F., and H.S. contributed data. A.J.H. performed all analyses. H.S. coordinated writing of the manuscript, with contributions from all authors.

## DATA AVAILABILITY

All new raw sequence data generated for this study have been deposited (or are in the process of being so) in the European Nucleotide Archive. Code and additional associated data will be made publicly available at https://github.com/ajhelmstetter/PAFTOL_magnoliids upon publication of this article.

## APPENDICES

### APPENDIX 1. Taxonomic treatment

**Meiocarpidieae Chatrou, Couvreur & Erkens, *tribus nov.*,** based on Meiocarpidioideae Chaowasku, Acta Bot. Brasil. 34(3): 525 (2020).

Type: *Meiocarpidium* Engl. & Diels

Note: See (Chaowasku, 2020) for details about morphological characterization of the tribe.

Mauloutchieae Ezedin & Sauquet, *tribus nov*.

Type: Mauloutchia Warb.

Description: Middle to high canopy trees or very rarely lianas (*Pycnanthus* spp.); dioecious or monoecious. **Sap** reddish to brownish or yellowish then oxidizing red. **Leaves** abaxially glaucous (*Cephalosphaera* and *Pycnanthus*) or not, papillose hairs absent or present and often early glabrescent, dots absent or rarely present (*Staudtia*); vernation convolute, lines sometimes present. **Inflorescences** densely globose cyme (*Staudtia*) or panicle (the rest).

**Flowers** unisexual; perianth 3–4(–5)-merous, green-yellow to orangish-red, spreading or reflexed at anthesis; synandrium stalk terete and convex, exserted or inserted but not completely enclosed by the perianth lobes (*Brochoneura* and *Doyleanthus*), unbranched monocyclic or branched (*Mauloutchia*), when branched with or without distinct phyllotaxy, anthers 3–5(–60), longifixed and fused or basifixed and free when branched. **Pollen** globose to boat shaped, aperture ulcerate or sulcate, exine sculpturing continuous to granulately verrucate-rugulate or minutely reticulate with psilate spines (*Pycnanthus*). **Fruits** dehiscent, globose to ellipsoid, glabrous or glabrescent, greenish to orangish-brown; pericarp thin (*Cephalosphaera*) to thick (*Pycnanthus*), smooth to slightly ridged or rarely carinate (*M. chapelieri*), leathery. **Arils** rudimentary to well developed or absent (*Mauloutchia*), when present partly enclosing the seed or rarely entire (*Staudtia*), deeply lacinate or only near apex (*Staudtia*), opaque creamy white (*Cephalosphaera*) to deep red (*Staudtia*). **Seeds** ellipsoid to ovoid; albumen not ruminate (except in *Pycnanthus*), containing oil.

Genera included: *Cephalosphaera* Warb. (1), *Doyleanthus* Sauquet (1), *Mauloutchia* Warb. (9), *Brochoneura* Warb. (3), *Pycnanthus* Warb. (4), and *Staudtia* Warb. (2–3).

Notes: This tribe displays the widest array of diversity across several traits, making it difficult to characterize morphologically. In particular, there is high diversity in anther morphology when compared to the remainder of the family. This is especially true of *Mauloutchia*, where the synandrium is branched, often with distinct phyllotaxy, and with basifixed anthers, which is unique in the family (Sauquet, 2004). The only other genus in which a non-entire synandria can be found is in a few species of *Knema* with lobed synandria.

Monoecy is widespread in the tribe, whereas dioecy appears randomly, specifically in *Brochoneura* and *Pycnanthus* which have been observed to contain individuals that are either dioecious or monoecious (Kühn and Kubitzki, 1993; Sauquet et al., 2003). Endosperm rumination is largely absent in this tribe, being only present in *Pycnanthus* (Kühn and Kubitzki, 1993), which would suggest ruminations as being derived in the family and having evolved more than once. *Doyleanthus* has yet to be molecularly sampled, however its morphology strongly suggests placement near the *Brochoneura*–*Mauloutchia* clade (Sauquet, 2003).

### Scyphocephalieae Ezedin & Sauquet, *tribus nov*

Type: Scyphocephalium Warb.

Description: High canopy trees; dioecious. **Sap** red. **Leaves** abaxially not glaucous, hairs rusty ferruginous and persistent, dots present; vernation convolute. **Inflorescences** densely globose cyme, sessile or pedunculate, in up to three heads, each head subtended by 2 bracts. **Flowers** unisexual; perianth (3–)4–5-merous, pinkish red, spreading at anthesis; synandrium stalk terete and flat-topped, inserted, unbranched monocyclic, anthers 6–10, fused longifixed. **Pollen** boat shaped, aperture ulcerate, exine sculpturing reticulate with muri subunits triangular and forming a crotonoid pattern. **Fruits** indehiscent, irregular globose, pubescent, orangish brown; pericarp fleshy. **Arils** entire. **Seeds** flattened; albumen ruminate, containing oil.

Genera included: *Scyphocephalium* Warb. (2).

Notes: Much remains unknown regarding the enigmatic *Scyphocephalium*. Possible synapomorphies include inflorescences pedunculate cymes bearing densely globose heads, pollen exine sculpturing in triangular crotonoid pattern, fruits indehiscent, and seeds flattened.

### Horsfieldieae Ezedin & Sauquet, *tribus nov*

Type: *Horsfieldia* Willd.

Description: Shrubs to mid canopy trees; dioecious or monoecious (*Endocomia*). **Sap** orange-reddish. **Leaves** abaxially glaucous or not, papillose hairs absent or rarely present (*H. iryaghedhi*), dots absent or rarely present (*Horsfieldia* spp.); vernation conduplicate.

**Inflorescences** paniculate with caducous basal cataphylls, bracts small or large and caducous, terminal buds absent. **Flowers** unisexual; perianth (2–)3–4(–5)-merous, green to orange, barely open or spreading to slightly recurved at anthesis; synandrium stalk terete and concave (rarely convex in *Endocomia*); anthers (2–)5–20(–30), fused longifixed; ovary ovoid (rarely ovoid-ellipsoid in *Horsfieldia*), glabrous or pubescent; stigma usually small, narrow or broad, 2-lobed, each lobe entire or (2–)3–5(–6)-lobulate. **Pollen** boat shaped, aperture sulcate, exine sculpturing often reticulate or rarely rugulate (*Gymnacranthera*). **Fruits** dehiscent, globose to ellipsoid, glabrescent to pubescent (rarely glabrous in *Horsfieldia* spp.), greenish yellow to dark orange; pericarp often thick leathery or somewhat fleshy, with or without tubercles, perianth sometimes persistent (*Horsfieldia*). **Arils** well developed, entire or shallowly (*Horsfieldia* spp.) to deeply (*Gymnacranthera*) lacinate, yellowish to red. **Seeds** ellipsoid, pointed at the apex (in *Endocomia*) or not, variegated (in *Endocomia*) or not; albumen ruminate, containing oil, starch absent; cotyledons divaricate, connate at base.

Genera included: *Endocomia* W.J.de Wilde (4), *Gymnacranthera* (A.DC.) Warb. (7), and *Horsfieldia* Willd. (106).

Notes: Possible synapomorphies for this clade include inflorescences subtended with basal cataphyll scars and seeds containing oil but no starch.

### Myristiceae Ezedin & Sauquet, *tribus nov*

Type: *Myristica* Gronov.

Description: Subcanopy to mid canopy trees; dioecious. **Sap** orangish-reddish or clear then oxidizing orange-red. **Leaves** abaxially glaucous or not, papillate hairs absent or sometimes present (*Myristica* spp.), dots present or absent; vernation conduplicate.

**Inflorescences** short branched paniculate or short wart-like, continuously growing woody brachyblasts from which flowers are successively borne, lacking basal cataphylls, bracteoles minute and caducous. **Flowers** unisexual; perianth (2–)3–4(–5)-merous, yellowish to reddish-brown, spreading and reflexed at anthesis; synandrium stalk terete and convex (*Myristica*) or concave (*Paramyristica*), flattened into a disk (*Knema*), inserted or rarely exserted (*Knema* spp.), unbranched monocyclic or rarely sublobate and whorled (i.e., *K. celebica*), anthers (3–)9– 25(–30), fused longifixed; ovary subglobose to oblong (rarely fusiform), pubescent; stigma often minutely 2-lobed, each lobe (2–)3–5(–6)-lobulate (in *Knema*). **Pollen** boat shaped (*Knema*) or elliptical and approaching boat shaped (*Myristica*), aperture sulcate, exine sculpturing reticulate to rugulate. **Fruits** dehiscent, globose to ellipsoid, glabrescent or pubescent (rarely glabrous in *Myristica* spp.), light greenish-yellow to orange; pericarp smooth, thick and fleshy (*Myristica*) or leathery (*Knema*). **Arils** well developed, lacinate completely or partly so to the base (*Myristica*) or entire and rarely lacinate at the apex only (*Knema*), yellow to deep red. **Seeds** ellipsoid, not variegated; albumen ruminate, containing oil and starch; cotyledons divaricate or rarely suberect (*Knema* spp.), connate at base and edges (*Myristica*) or very slightly to rarely so (*Knema*).

Genera included: *Knema* Lour. (96), *Myristica* Gronov. (172), and *Paramyristica* de Wilde (1). Notes: The close relation between *Myristica* and *Knema* is well supported molecularly.

Although dioecious, a few individuals of *Myristica* have been reported as monoecious by de Wilde (1984) who considered these cases anomalous. *Paramyristica* is tentatively included here based on morphological grounds and partly from the results of our phylogenetic analyses, however its topological placement remains unstable. Possible synapomorphies for this clade are inflorescences lacking basal cataphyll scars and seeds containing oil and starch. However, *Paramyristica* notably deviates from the tribe as it has cataphyll scars and lacks bracteoles (de Wilde, 1994).

### Viroleae S.M.Oliveira & Sauquet, *tribus nov*

Type: *Virola* Aubl.

Description: Mid to high canopy trees or very rarely shrubs (*Virola sessilis* and *V. subsessilis*); dioecious or rarely monoecious (*Iryanthera* spp.). **Sap** red, brownish, pinkish, sometimes initially watery then oxidizing opaque white (*Otoba* spp.) or straw-coloured (*Osteophloeum*). **Leaves** with persistent indument or glabrescent; hairs sessile stellate, dendritic, stipitate stellate or malpighiaceous (*Iryanthera* spp.); vernation convolute or conduplicate (*Otoba*), lines sometimes present. **Inflorescences** axillary, supra axillary or less often cauliflorous (*Iryanthera* spp.), racemose or variously branched paniculate; bracts mostly caduceus; bracteoles present or not. **Flowers** unisexual; perianth 3–4(–5)-merous, 1-4 cm long, partly connate, often fleshy, yellow or slightly green to whitish; synandrium stalk terete, flat to convex; anthers 2–10(–20), fused longifixed or free basifixed (in *Compsoneura* spp., *Otoba* spp.), dehiscing by longitudinal slits; ovary sessile or short stipitate, globose, subglobose, ovoid, ellipsoid or pyriform, glabrous or pubescent, unicarpellate, unilocular; stigma sessile, 2-lobed.

**Pollen** in monads or exceptionally tetrads (*Iryanthera*); size 21–40 µm; mostly sulcate with sharply delimited aperture or sulcoidate with aperture not sharply defined; boat shaped or globose spherical; exine sculpturing reticulate or psilate (*Otoba*). **Fruits** dehiscent, 2-valvate, globose, subglobose or ellipsoid, glabrous, glabrescent or pubescent. **Arils** partly to completely enclosing the seed, red, opaque orangish-red or translucent white (*Otoba* spp.). **Seeds** ellipsoid, variegated (in *Compsoneura*) or not; albumen ruminate, containing oil; cotyledons often vestigial, divergent, and becoming haustorial during germination.

Genera included: *Bicuiba* W.J.de Wilde (1), *Compsoneura* (DC.) Warb. (17), *Iryanthera* (A.DC.) Warb. (22), *Osteophloeum* Warb. (1), *Otoba* (DC.) H.Karst. (12), and *Virola* Aubl. (71).

Notes: Outside of Mauloutchieae, this tribe contains the only other two genera which contain species that exhibit basifixed and free anthers: *Compsoneura* and *Otoba*.

## SUPPORTING INFORMATION

**APPENDIX S1.** Full voucher and source information for all taxa included in this study.

**APPENDIX S2.**
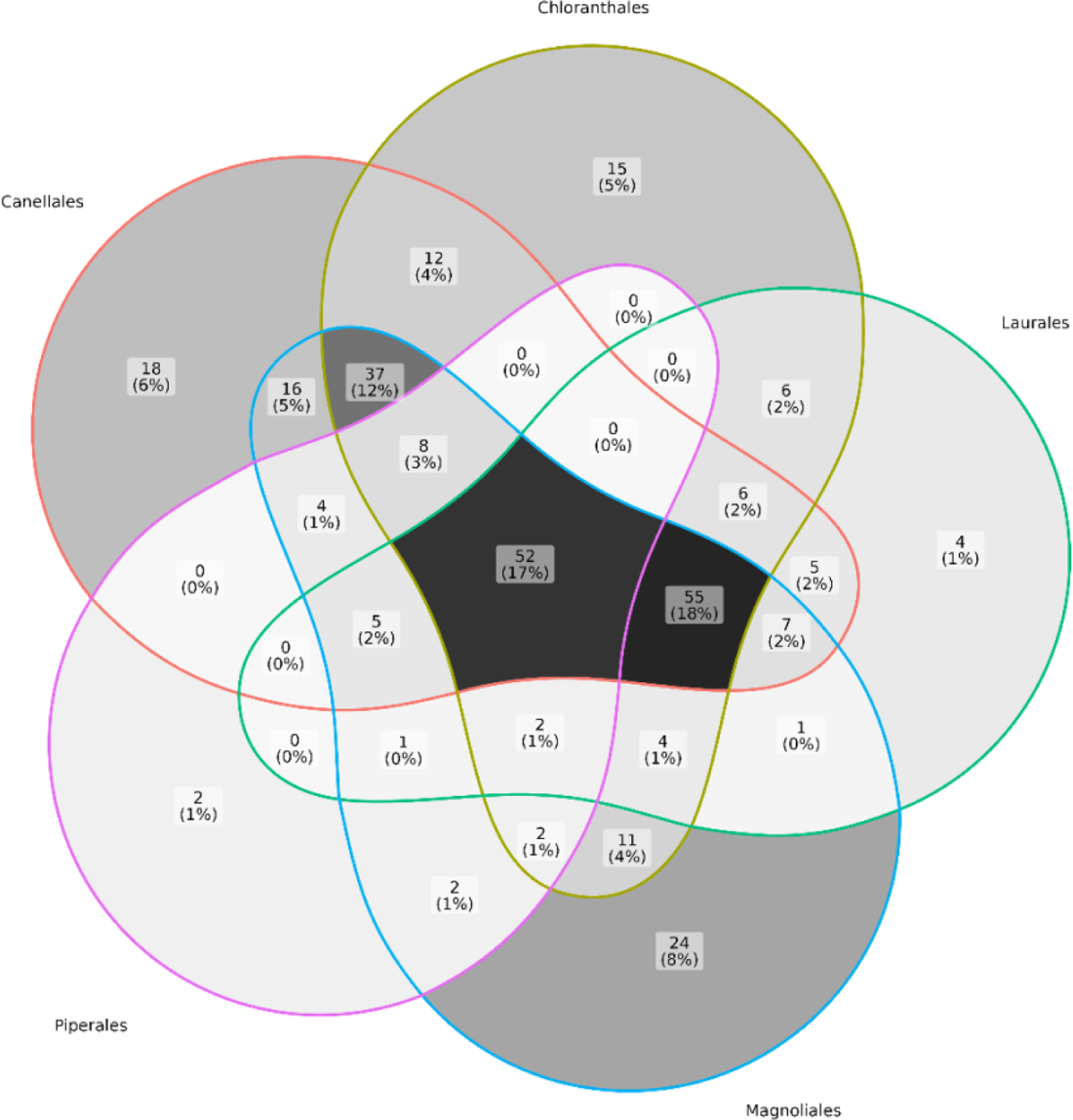
Venn diagram showing how loci that are selected as suitable for phylogenetic inference for each order are shared among orders. Filtering was conducted on each order separately to generate the set of loci for which at least 50% of the exon was recovered in at least 50% of specimens. Within each section of the Venn diagram, numbers indicate count of loci falling into the section and percentage of the total dataset (all sets from each order combined). The central section indicates that there are 52 loci (17% of the whole dataset) that are recovered well enough to be selected as suitable for phylogenetic inference in each of the five orders. The colored outer section of each order (e.g. red outline for Canellales) on the diagram indicates the number of loci that were selected as suitable only for a given order (e.g. 18 loci, or 6% of the whole dataset, for Canellales).

**APPENDIX S3.**
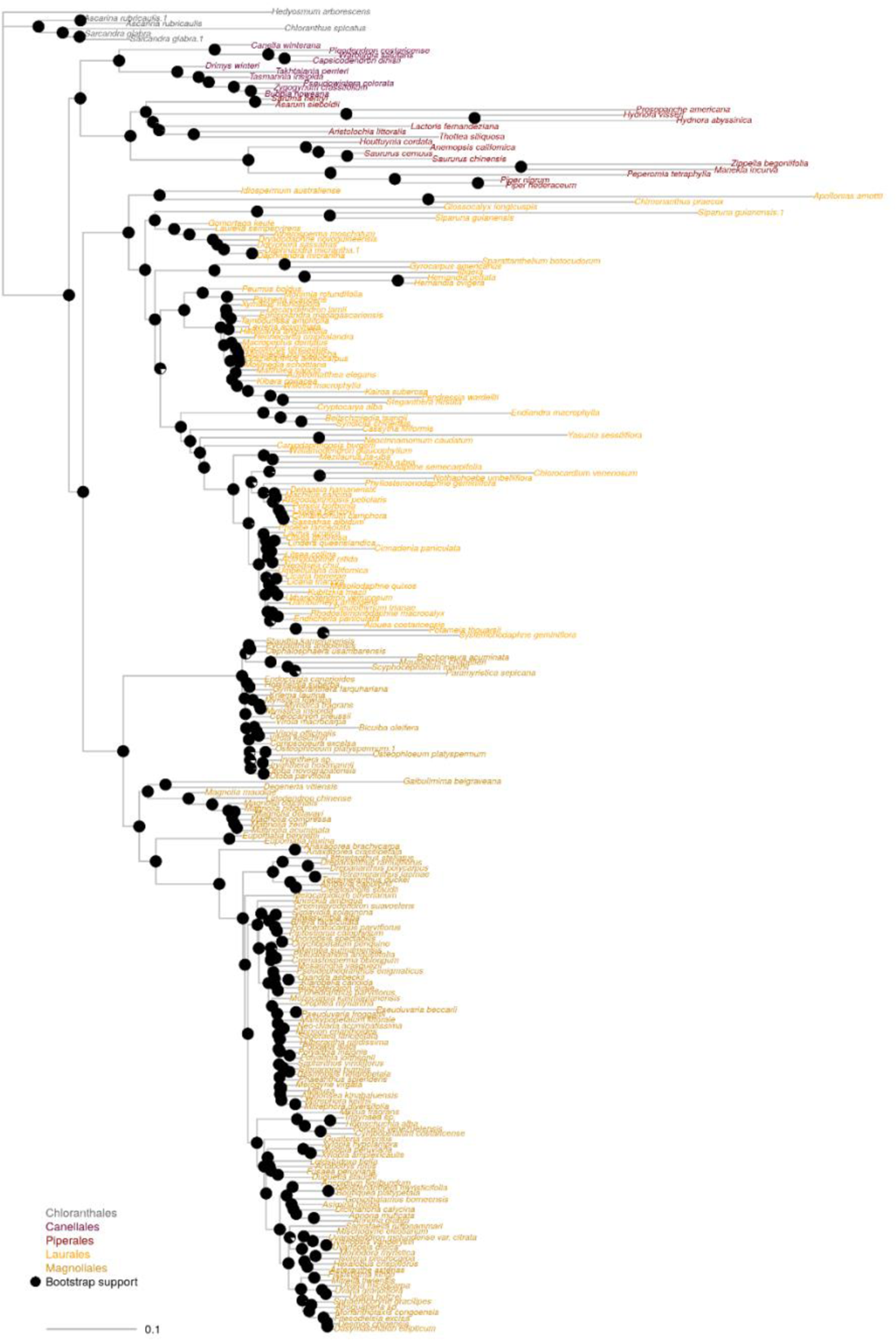
Phylogenetic relationships in Magnoliidae obtained in our main concatenation analysis. Pie charts at nodes represent bootstrap support values. Branches are in units of substitutions per site with a scale bar in the bottom left.

**APPENDIX S4.**
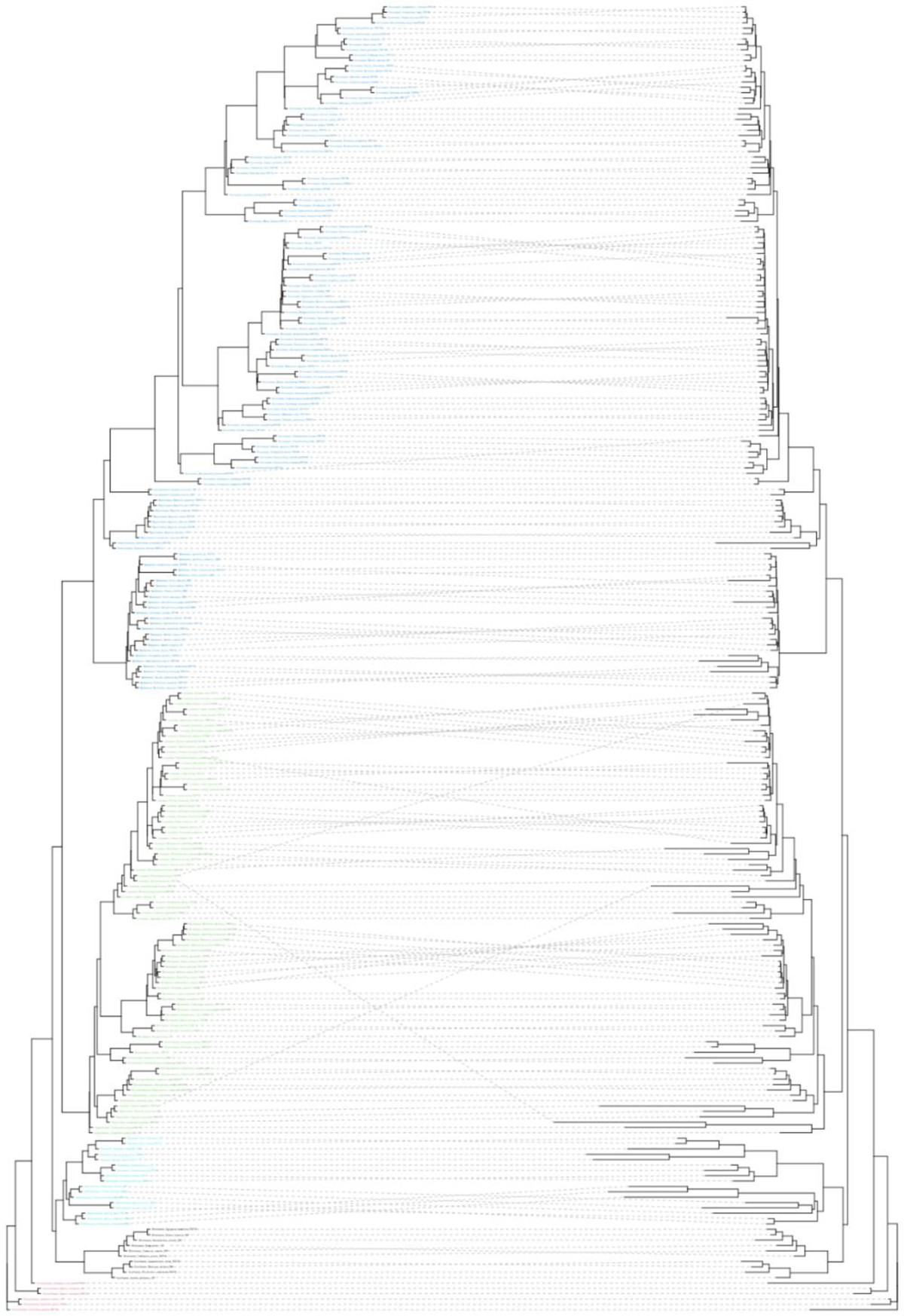
Tanglegram detailing topological differences between the main coalescent (Figs. 4, 5) and concatenation (Appendix S3) analyses. Grey dashed lines join the corresponding tips in each tree.

**APPENDIX S5.**
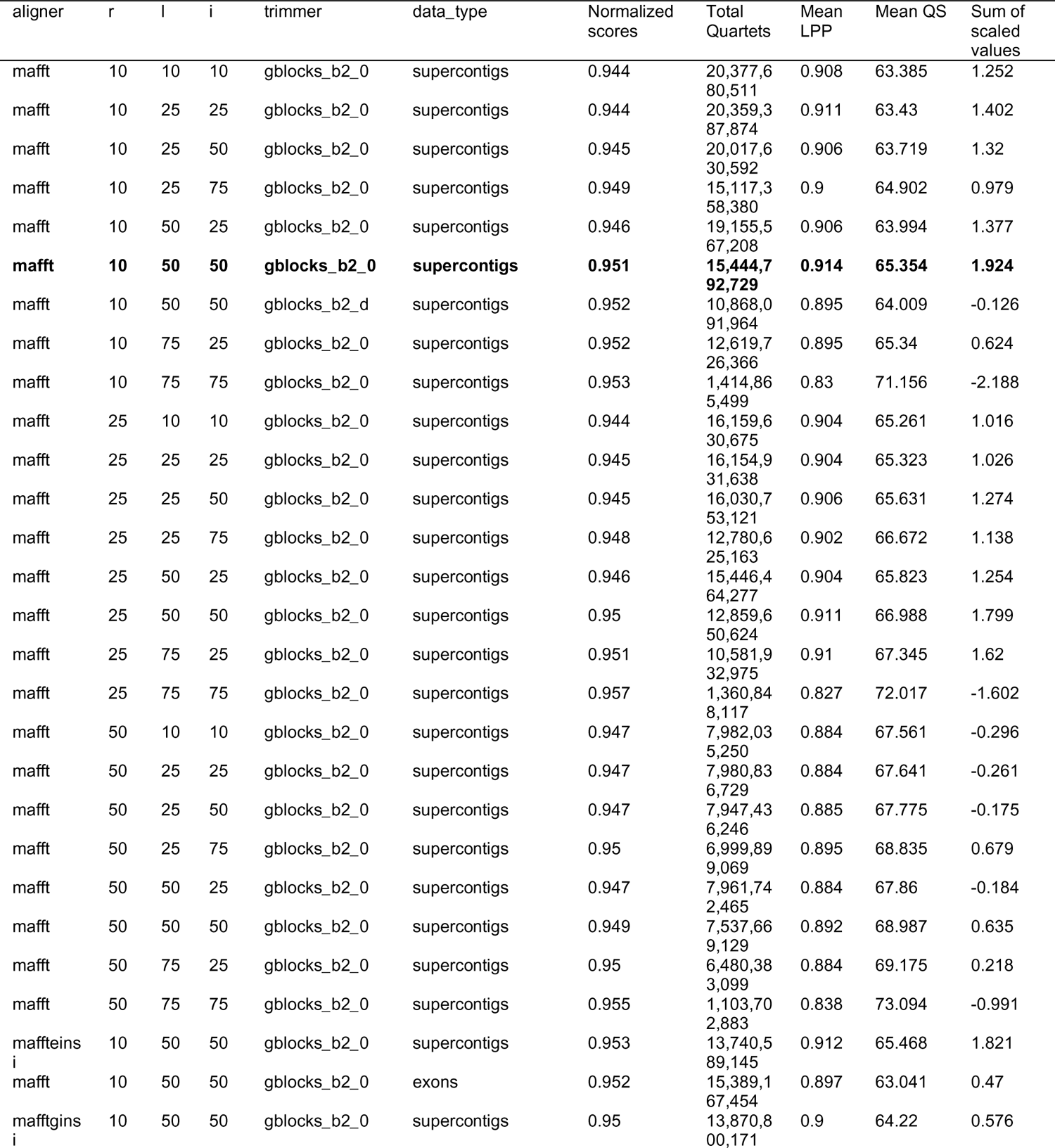
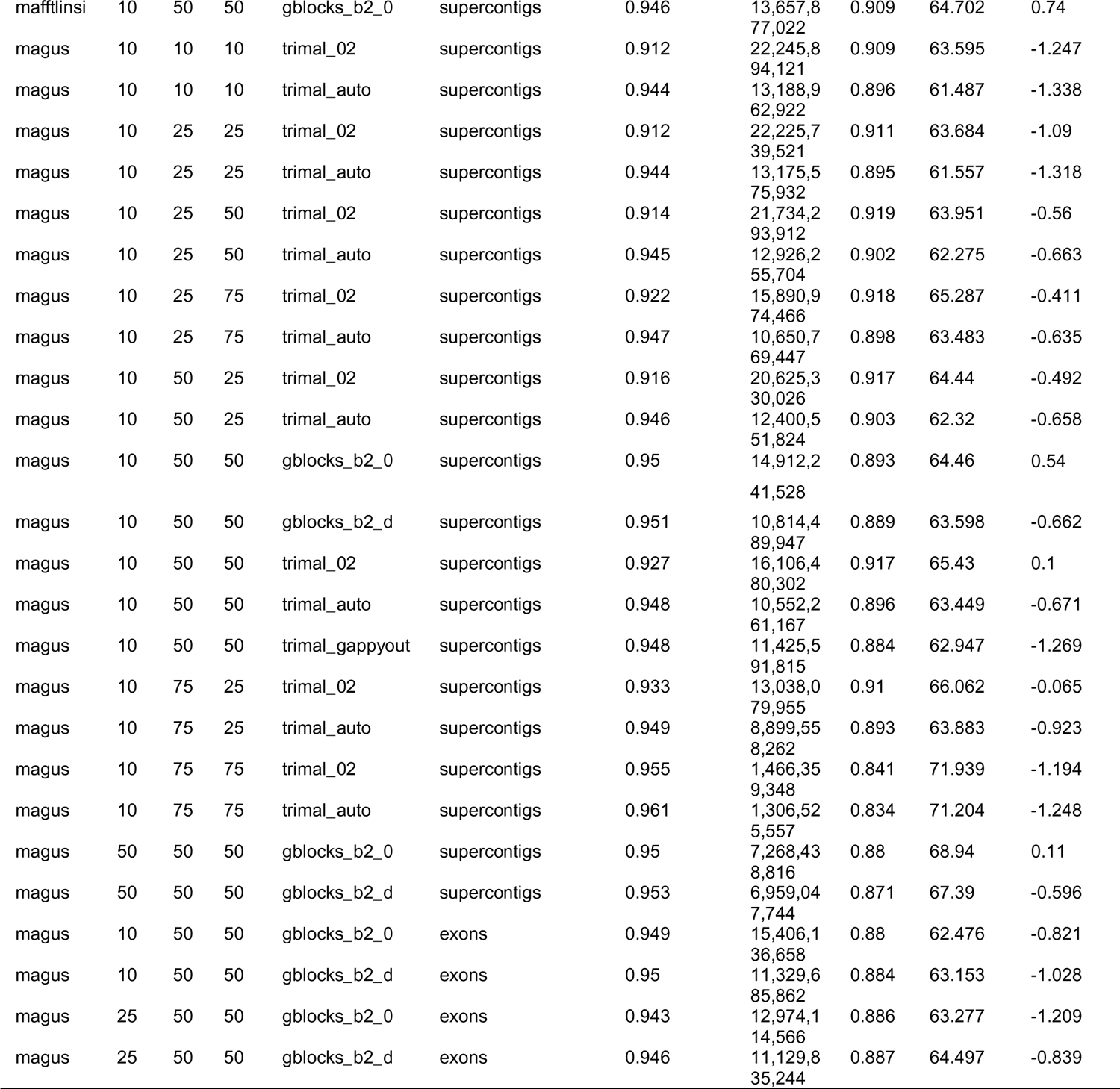
Table of mean quality scores per coalescent analysis using four different metrics outlined in the methods. The different characteristics of each analysis are shown in the first six columns. The final column is the sum of each row after the values were scaled. The analysis highlighted in boldface received the highest sum of scaled values across four quality metrics and was used as our main reference analysis presented in this paper (Figs. 4, 5).

**APPENDIX S6.**
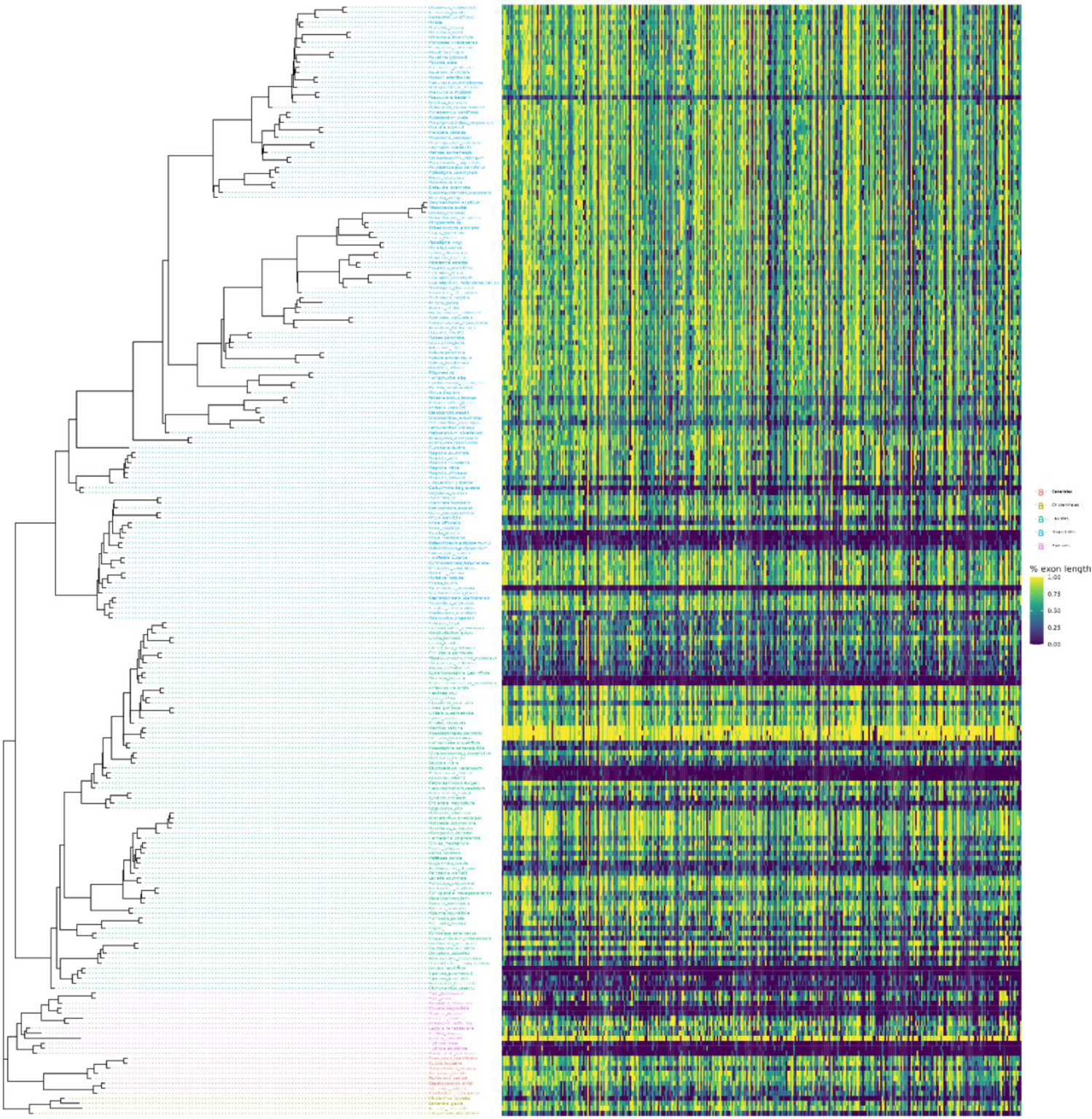
Coalescent tree from Figure 5 alongside a heatmap showing the proportion of exon recovered for all 353 exons in the Angiosperms353 kit.

**APPENDIX S7.**
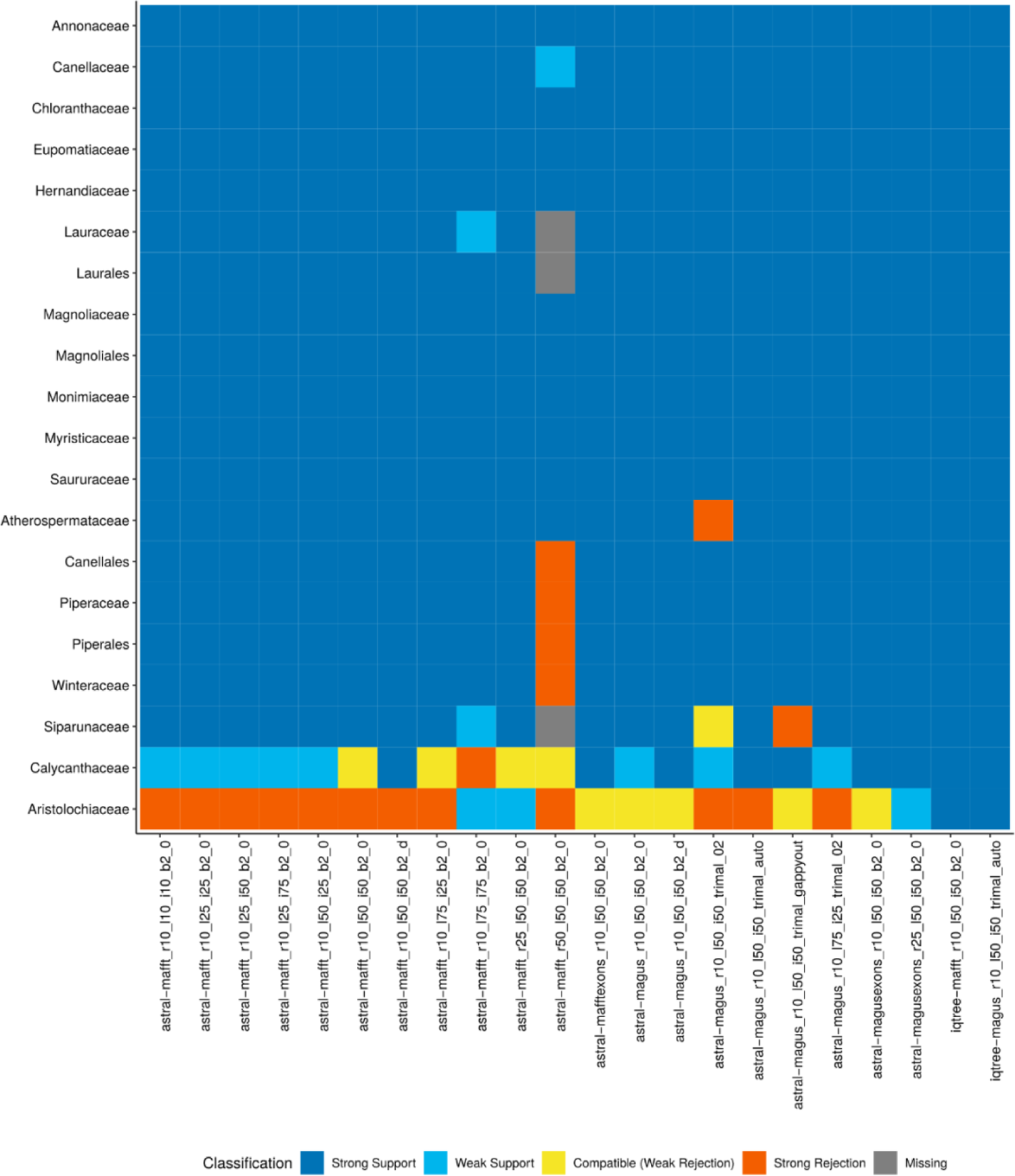
Heatmap showing how well supported major clades (families, orders) are across coalescent analyses from a range of different analytical approaches. Strong/weak support indicates whether the relevant branch of the gene tree had a local posterior probability (LPP) support value above/below a threshold of 0.9. Weakly rejected clades correspond to clades that are not present in the tree, but are compatible if low support branches (below 0.9 LPP) are contracted. If they are still not compatible they count as strongly rejected clades.

**APPENDIX S8.**
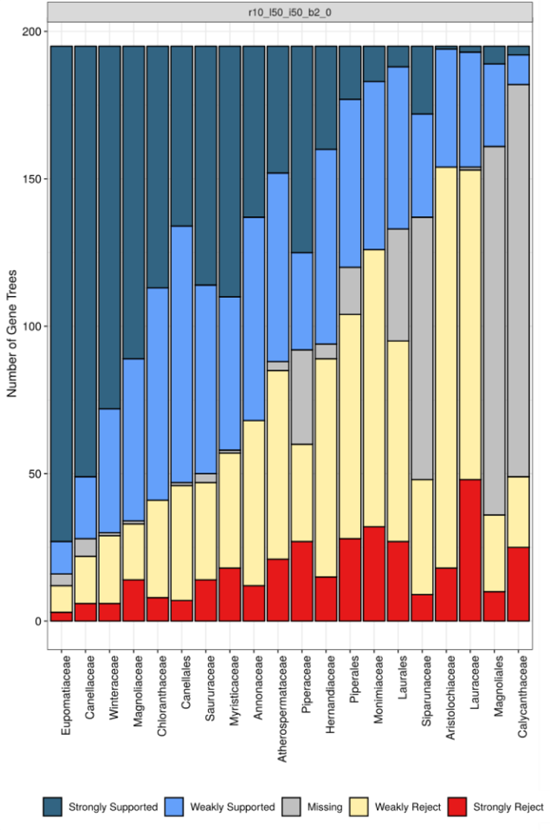
Stacked bar plots showing the number of gene trees that support the monophyly of major magnoliid clades (as in Appendix S7). The set of gene trees used were the same as those used for the tree in Figure 5. Strong/weak support indicates whether the relevant branch of the gene tree had a bootstrap support value above/below a threshold of 90%. Weakly rejected clades are those that are not in the tree but are compatible if low support branches (below 75%) are contracted. Those branches that were not compatible after contracting low support branches count as a strong rejection.

**APPENDIX S9.**
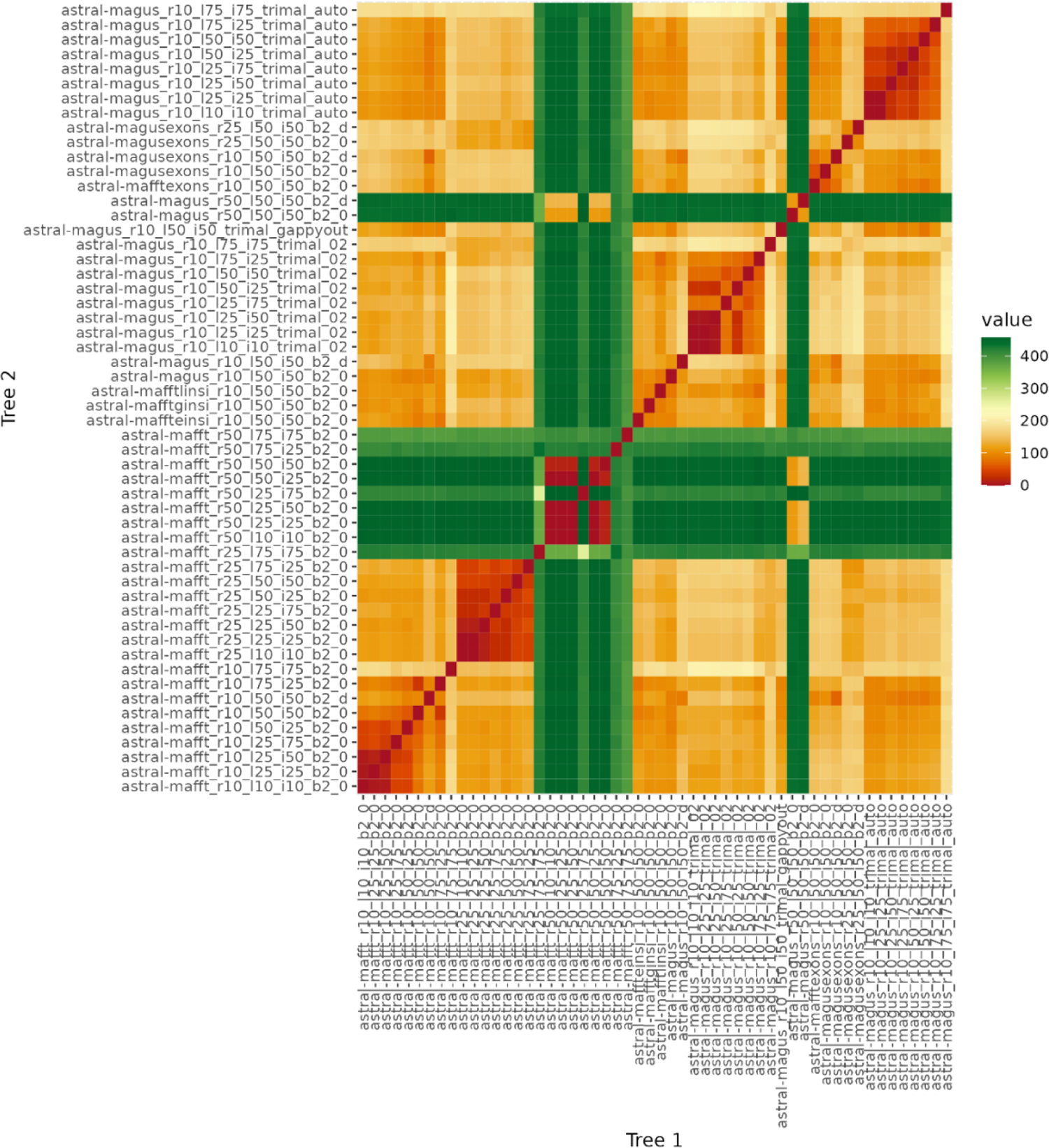
Heatmap depicting the Robinson-Foulds distances between each pair of trees generated by our analyses.

**APPENDIX S10.**
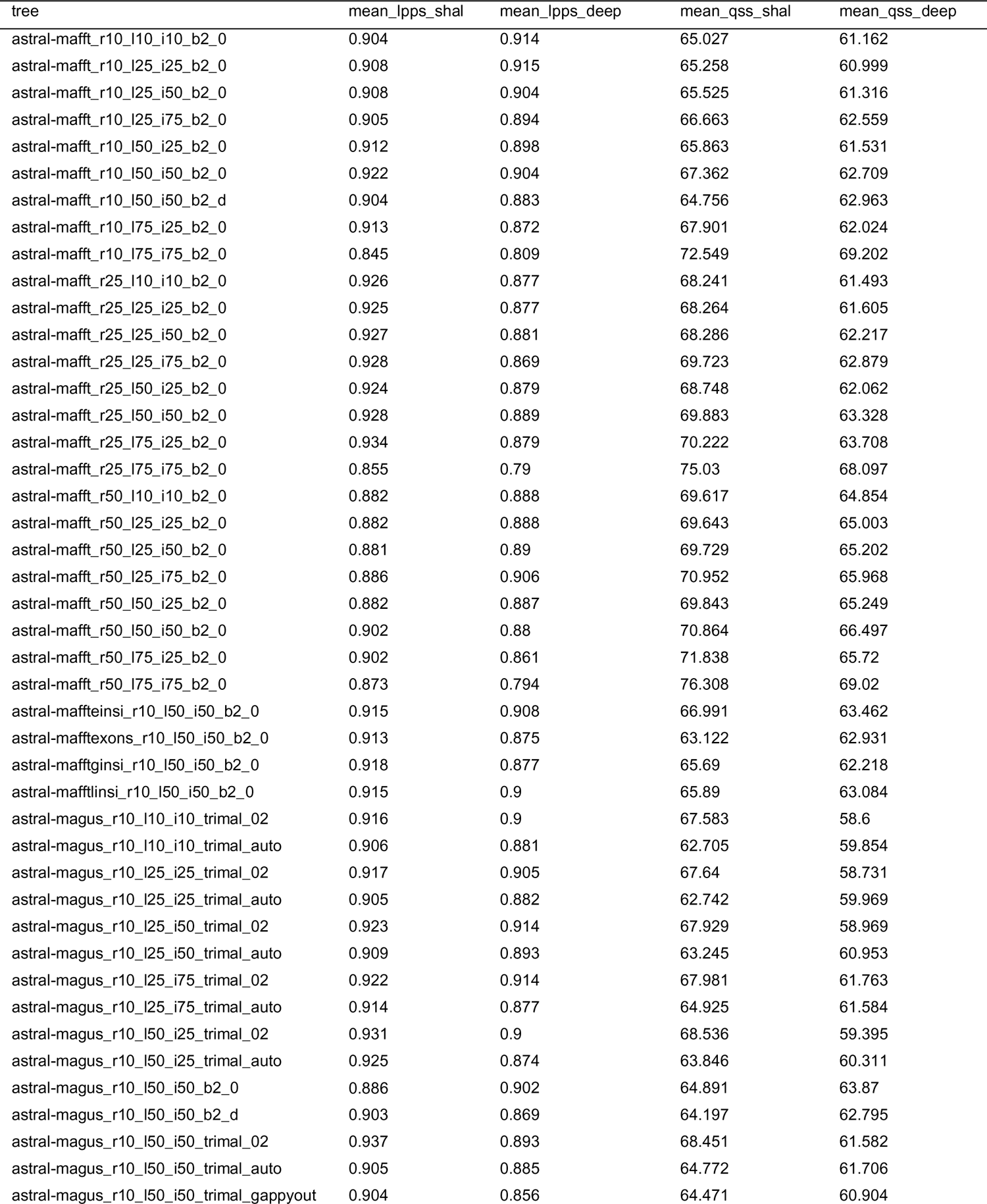
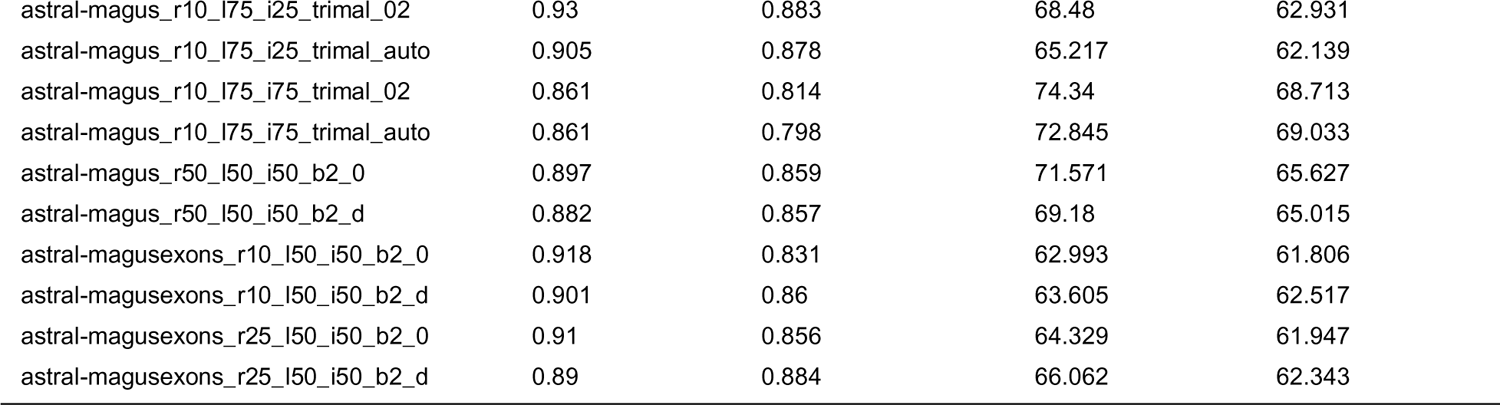
Table of mean Local Posterior Probability (LPP) and Quartet Support (QS) scores per (ASTRAL) tree for shallow (depth < 5) and deep (depth >= 5) nodes.

**APPENDIX S11.**
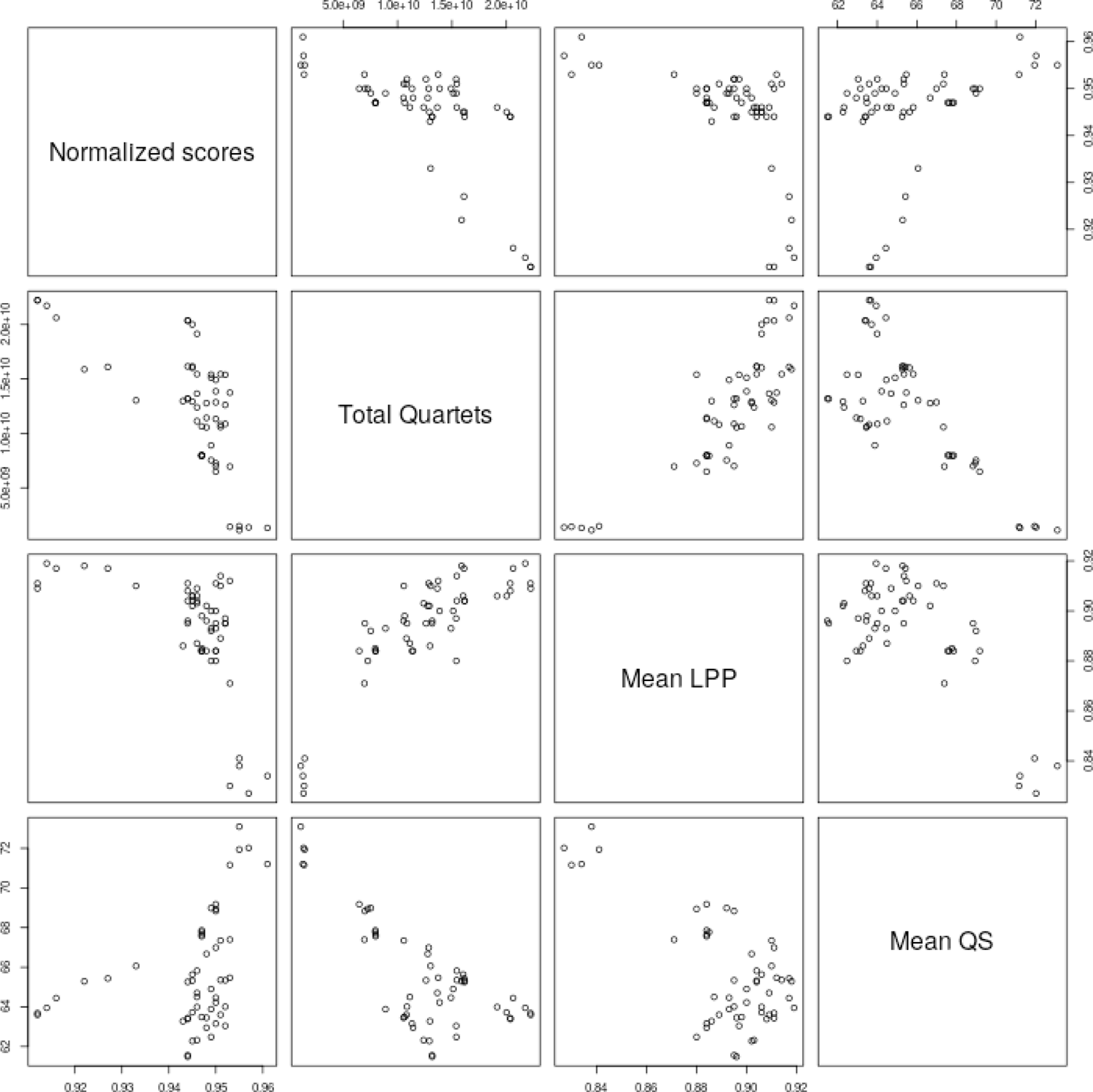
Pairwise scatterplots of the four different quality metrics used in our study. Each point represents an ASTRAL tree (as listed in Appendix S5).

**APPENDIX S12.** Phylogenetic trees obtained with the coalescent analyses conducted with different thresholds for locus filtering or different alignment strategies than the main reference analysis presented in Figs. 4 and 5.

